# Neural and cognitive correlates of performance in dynamic multi-modal settings

**DOI:** 10.1101/2022.03.23.485424

**Authors:** Chloe A. Dziego, Ina Bornkessel-Schlesewsky, Sophie Jano, Alex Chatburn, Matthias Schlesewsky, Maarten A. Immink, Ruchi Sinha, Jessica Irons, Megan Schmitt, Steph Chen, Zachariah R. Cross

## Abstract

The endeavour to understand human cognition has largely relied upon investigation of task-related brain activity. However, resting-state brain activity can also offer insights into individual information processing and performance capabilities. Previous research has identified electroencephalographic resting-state characteristics (most prominently: the individual alpha frequency; IAF) that predict cognitive function. However, it has largely overlooked a second component of electrophysiological signals: aperiodic 1/*f* activity. The current study examined how both oscillatory and aperiodic resting-state EEG measures, alongside traditional cognitive tests, can predict performance in a dynamic and complex, semi-naturalistic cognitive task. Participants’ resting-state EEG was recorded prior to engaging in a Target Motion Analysis (TMA) task in a simulated submarine control room environment (CRUSE), which required participants to integrate dynamically changing information over time. We demonstrated that the relationship between IAF and cognitive performance extends from simple cognitive tasks (e.g., digit span) to complex, dynamic measures of information processing. Further, our results showed that individual 1/*f* parameters (slope and intercept) differentially predicted performance across practice and testing sessions, whereby flatter slopes were associated with improved performance during learning, while higher intercepts were linked to better performance during testing. In addition to the EEG predictors, we demonstrate a link between cognitive skills most closely related to the TMA task (i.e., spatial imagery) and subsequent performance. Overall, the current study highlights (1) how resting-state metrics – both oscillatory and aperiodic - have the potential to index higher-order cognitive capacity, while (2) emphasising the importance of examining these electrophysiological components within more dynamic settings and over time.

## 1. INTRODUCTION

The endeavour to understand the relationship between neural activity and cognition typically relies upon investigating on-task (or evoked) neural dynamics. However, emerging research demonstrates that resting-state brain activity offers invaluable insight into individual differences in information processing and performance capacities (Grandy, Werkle-Bergner, Chicherio, Lövdén, et al., 2013; Klimesch, 1999; Ouyang et al., 2020; Voytek et al., 2015). Hans Berger first proposed in 1929 that, “mental work … adds only a small increment to the cortical work which is going on continuously and not only in the waking state” (as cited in Snyder & Raichle, 2012, p. 3). This claim is based on the observation that the human brain consistently demonstrates activity independent of external stimulation (Snyder & Raichle, 2012), likely reflecting intrinsic properties of its internal networks (i.e., underlying anatomical structure, local neuronal dynamics, signal transmission delays and noise; Deco et al., 2011). From this perspective, resting-state neural activity may provide insight into the global functional capacity of the brain by reflecting homeostatic mechanisms (Marder & Goaillard, 2006) that facilitate or impede optimal cognitive function during task engagement. Investigation of brain activity across neurological and psychiatric disorders, as well as in ageing, has demonstrated how adverse changes in cognition and behaviour can be paralleled by distinct resting-state profiles (e.g., Damoiseaux et al., 2008; Greicius, 2008; Klimesch, 1999; Newson & Thiagarajan, 2019; Voytek et al., 2015). However, while these findings demonstrate how resting-state activity relates to dysfunction, its role in allowing for peak cognitive functioning during dynamic information processing is considerably less well understood. As resting-state activity is thought to provide an indication of the brain’s functional capacity (Deco et al., 2011), it may offer insights into neurophysiological characteristics that facilitate effective information processing and performance.

While multiple neuroscientific methods have been used to investigate resting-state dynamics (Biswal, 1995; Klimesch, 1999; Raichle et al., 2001), EEG is one of the most widely used techniques to examine the relationship between cognitive functions and brain activity (Bastiaansen et al., 2011). The continuous EEG signal is comprised of two components: periodic (oscillatory activity) and aperiodic (scale-free/fractal) 1/*f* activity (Buzsáki, 2006; Wen & Liu, 2016). The oscillatory component encompasses well-defined frequency bands (i.e., alpha, beta, theta, delta, gamma) that demonstrate intrinsic respective time-scales (Kayser & Ermentrout, 2010). In the exploration of individual differences in cognitive function and resting-state EEG activity, the alpha band (~ 7 – 13 Hz; Klimesch, 1999) has been of particular interest, with the individual alpha frequency (IAF) hypothesised to be a physiological measure of cognitive ability (Grandy, Werkle-Bergner, Chicherio, Lövdén, et al., 2013; Grandy, Werkle-Bergner, Chicherio, Schmiedek, et al., 2013). The IAF refers to the peak frequency within the alpha band (Klimesch, 1999) and is a property of the human EEG that exhibits trait-like characteristics (Grandy, Werkle-Bergner, Chicherio, Schmiedek, et al., 2013). IAF has been found to positively correlate with IQ (Grandy, Werkle-Bergner, Chicherio, Lövdén, et al., 2013), memory performance (Klimesch, 1996, 1999), visuo-perceptual ability (Cecere et al., 2015; Samaha & Postle, 2015) and memory outcomes following sleep (Chatburn et al., 2021; Cross, Zou-Williams, Wilkinson, Schlesewsky & Bornkessel-Schlesewsky, 2020). However, the relationship between alpha and cognitive performance is not always reliably exhibited (Ouyang et al., 2020). For example, a U-shaped relationship has been observed between alpha and somatosensory perception (Lange et al., 2014; Linkenkaer-Hansen et al., 2004; Zhang & Ding, 2009) and individuals with lower IAF demonstrate superior performance in spatial localisation tasks (Howard et al., 2017). In linguistic contexts, differences in IAF have been proposed to reflect distinct information processing strategies, with lower/higher IAF not more advantageous than the other (Bornkessel et al., 2004; Kurthen et al., 2020). These findings demonstrate the complexities in the relationship between alpha and cognition and could suggest that other electrophysiological phenomena may be interacting with or conflating these findings (Cross, Corcoran, Schlesewsky, Kohler & Bornkessel-Schlesewsky, 2020; Donoghue et al., 2020, 2021).

The second component of EEG (aperiodic 1/*f* activity) can be observed when the EEG signal is transformed into its power spectral density (for a schematic of oscillatory vs aperiodic signal decomposition, see Figure 1). Oscillatory activity is observed through multiple peaks at specific frequencies, with the aperiodic component manifesting as a distinguishable descending quasi-straight line (Miller et al., 2009). This line subscribes to the 1/*f* power-law relationship of *S*(*f*) ∝ 1/*f*^*β*^ (He, 2014), whereby lower frequencies exhibit greater amplitudes (i.e., power), while higher frequencies display decreased amplitude (Kayser & Ermentrout, 2010). The power-law exponent (*β*) represents the steepness of the 1/*f* slope (He, 2014). A *β* value of 0 reflects random variation in power or ‘white noise’ – demonstrated through a horizontal (or flat) plotted line in the power spectrum density (PSD) plot – while increasing *β* values indicate increasing systematic variation (He, 2014). In young healthy adults, scalp electrophysiology typically demonstrates an exponent (*β*) between 1 and 2 (He, 2014). This activity is referred to as aperiodic, as it is not confined to a specific time scale (Wen & Liu, 2016). In traditional electrophysiological analyses, 1/*f* activity is typically ignored or treated as a nuisance variable (for a detailed discussion, see Donoghue et al., 2020; He, 2014; Lendner et al., 2020), with a considerable portion of extant literature largely focusing on oscillatory activity. However, changes in the aperiodic exponent and intercept can be misinterpreted as increases or decreases in oscillatory power, potentially leading to the disparity in previous findings (for detailed discussion, see Donoghue et al., 2020). It is thus important to disentangle the influence of aperiodic 1/*f* activity and neural oscillations to gain a better understanding of the neurobiological basis of cognition in health, ageing and disease (Wen & Liu, 2016).

**Figure 1.**
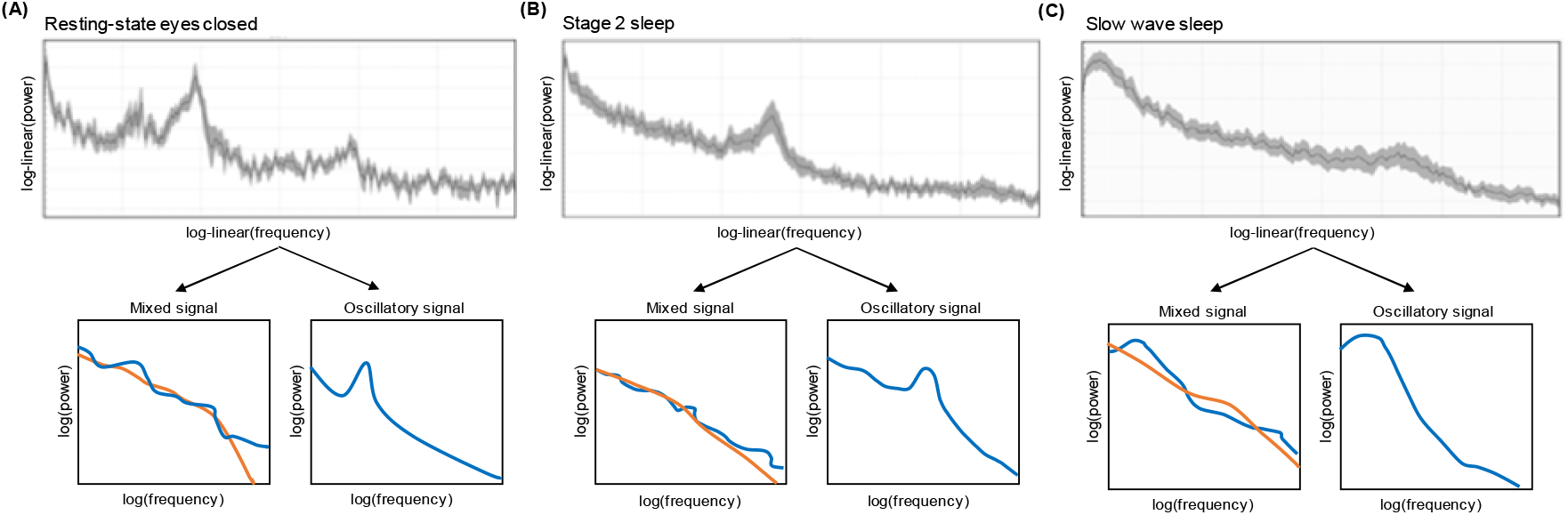
Schematic of power spectral density (PSD) estimates of fractal (aperiodic 1/*f*) and oscillatory activity across different states of arousal. **(A)** PSD plot from 30s of an eyes-closed resting state period. **(B)** PSD plot from 30s of stage 2 sleep. **(C)** PSD plot from 30s of slow wave sleep. The top row illustrates the raw EEG, while the bottom row illustrates the mixed (left) and separated (right) oscillatory signal independent from the background, aperiodic signal (orange line). Log transformed power (i.e., strength of neural activity) is represented on the y-axis (higher values indicate greater activity), while frequency (Hz; cycles per second) is represented on the x-axis (values further to the right indicate higher frequency). Note the clear spectral peak in the 10 Hz range in **(A)**, which reflects the peak alpha frequency.

Beyond the basic physiological basis of the 1/*f* exponent, research is beginning to demonstrate the functional relevance of aperiodic activity in human cognition (Cross, Corcoran, et al., 2020; Dave et al., 2018; Donoghue et al., 2020; Immink et al., 2021; Ouyang et al., 2020). For example, slower processing speed, reflected by longer reaction times in an object recognition task, has been associated with a flatter resting-state 1/*f* slope (Ouyang et al., 2020), while superior performance on a visual working memory task is related to a steeper 1/*f* slope (Donoghue et al., 2020). A steeper 1/*f* slope and a higher intercept (thought to reflect higher mean neural population spiking; Manning et al., 2009; Miller et al., 2012) have also been associated with higher perceptual sensitivity in a visuomotor learning task (Immink et al., 2021) as well as learning complex rules in an artificial language learning paradigm (Cross, Corcoran, et al., 2020). Research in this area has also reported that resting aperiodic activity predicts performance over and above that of resting alpha power (Donoghue et al., 2020; Ouyang et al., 2020), attesting to the importance of further investigation of these components in conjunction.

In support of the relationship between aperiodic activity and cognitive function, 1/*f* slopes have additionally been related to changes in cognition that parallel ageing. The resting aperiodic slope becomes flatter in elderly populations (Dave et al., 2018; Donoghue et al., 2020; Voytek et al., 2015) and is associated with declines in cognitive performance, as demonstrated in working memory tasks (Donoghue et al., 2020; Voytek et al., 2015). Further, in a study of language prediction in younger and older adults, flatter 1/*f* slopes were related to decreased N400 amplitudes, thought to index reduced strengths of predictions in anticipatory language processes (Dave et al., 2018). Taken together, this emerging evidence demonstrates the strong predictive capacity of resting-state-derived aperiodic activity in determining cognition across a range of modalities, developmental stages, and disease states.

However, despite the associations between aperiodic activity and higher-order cognition, there is little consensus regarding the exact mechanisms underlying the 1/*f* slope and intercept. The intercept is posited to reflect mean neural population spiking (i.e., overall power; Donoghue et al., 2020; Manning et al., 2009; Miller et al., 2012) and has been related to the fMRI BOLD signal (Winawer et al., 2013). However, both an increase (Immink et al., 2021) and decrease (Ouyang et al., 2020) in the 1/*f* intercept has been related to improved visuomotor and cognitive performance, suggesting that further research is necessary to disentangle these relationships. Moreover, while steeper resting-state-derived 1/*f* slopes seem to be related to improved cognitive performance, there is considerably less evidence linking the 1/*f* intercept to behavioural and cognitive outcomes (cf. Cross, Corcoran, et al., 2020; Immink et al., 2021).

The largest body of evidence proposes that aperiodic activity, particularly the 1/*f* slope, reflects the excitation/inhibition balance within recurrent neural networks (Destexhe et al., 2001; Gao et al., 2017). This concept stems from initial findings suggesting aperiodic activity reflects neuronal firing rates (Buzsáki et al., 2012; Miller et al., 2009), with greater asynchronous activity related to flatter PSD. Experimental manipulation supports this perspective: alcohol (Stock et al., 2020) and propofol intake (Colombo et al., 2019; Lendner et al., 2020; Medel et al., 2020) – which are known to increase inhibitory GABAergic activity – are associated with steeper 1/*f* slopes. Individuals also exhibit steeper 1/*f* slopes during slow wave and rapid eye-movement sleep (states with increased inhibition), in comparison to their resting but awake states (Allegrini et al., 2015; Lendner et al., 2020). Those with disorders related to imbalances in excitation/inhibition, such as schizophrenia (Molina et al., 2020; Peterson et al., 2017; Slezin et al., 2007), mania (Bahrami et al., 2005), ADHD (Ostlund et al., 2021; Robertson et al., 2019) and Down Syndrome (Hemmati et al., 2013) also show altered 1/*f* slopes when compared to healthy controls. It has been suggested that the cognitive processing enhancements linked to steeper slopes may arise from an optimal balance between excitation and inhibition in the brain (Lendner et al., 2020; Weber et al., 2020), allowing greater consistency of stimulus processing and enhanced perceptual sensitivity (Immink et al., 2021; Tran et al., 2020).

Taken together, current experimental evidence demonstrates that performance on a variety of cognitive measures is associated with both oscillatory (e.g., IAF) and aperiodic (e.g., 1/*f* slope and intercept) resting-state-derived parameters, and that these associations differ across development and within disease states. However, research is yet to examine the association between intrinsic oscillatory and aperiodic activity and fluctuations in cognitive processing on more naturalistic tasks over-time. Here, we examined how individual resting-state EEG activity – both oscillatory and aperiodic – relate to performance on a dynamic multi-modal cognitive task known as Target Motion Analysis (TMA) within the Control Room Use Simulation Environment (CRUSE; developed by Defence Science and Technology Group, see Michailovs et al., 2021). CRUSE is a medium-fidelity submarine control room simulation comprising multiple interoperable stations, including sensors (e.g. SONAR, optronics), TMA and Track Management. It was developed to enable applied individual and team experimentation on psychological factors relevant to control room operations, such as situational awareness, workload, team processes and team performance. The primary task in CRUSE is building a picture of vessels (referred to as contacts) surrounding the submarine (i.e., creating a ‘tactical picture’). The role of the TMA operator involves integrating information from other operators (e.g. SONAR, Optronics and the Track Manager) to derive ‘solutions’ for each of the contacts within the submarine’s sensor range. Performance on the CRUSE task can be measured by both the speed and accuracy of plotted solutions, ability to integrate both visual and auditory information and the prioritisation of tracking various stimuli in a virtual environment. Performance is also tracked throughout the task in twenty second epochs, which allowed us to relate individual differences in resting-state neural activity to dynamic changes in behaviour over time. As such, by using a complex, semi-naturalistic task – in contrast to the typically implemented one-dimensional cognitive tests – we aim to further characterise the functional significance of resting-state-derived oscillatory and aperiodic neural activity in higher-order information processing.

We also obtained measurements on a range of traditional cognitive tasks, including measures of working memory, mental rotation, and statistical learning ability, with the aim of establishing how oscillatory- and aperiodic-based parameters explain dynamic task performance alongside that of traditional behavioural metrics. The evidence comparing oscillatory and aperiodic measures on higher-order information processing with more traditional cognitive measures remains scarce. The transferability of specific cognitive abilities to near and far domains has been heavily researched and debated amongst cognitive psychologists (Perkins & Salomon, 1989; Sala & Gobet, 2017). However, there is little evidence that greater performance in one area of cognition will be paralleled by greater performance across multiple other cognitive domains (Sala & Gobet, 2017) and reviews have found that training of one particular cognitive skill does not enhance performance on distantly related tasks or general cognition at all (Simons et al., 2016). In a similar fashion, we might predict that performance on traditional cognitive tasks (i.e., digit span) may not be associated with performance in a more complex and dynamic setting. Nevertheless, some cognitive skills are thought to be more transferable than others (e.g., spatial imagery; Uttal et al., 2013) and may act as stronger predictors of performance across a variety of settings. Including a selection of cognitive tests in the present study allowed us to examine both whether these would show a relationship with performance on the dynamic, naturalistic task under consideration and how they relate to the resting-state EEG metrics of interest.

Additionally, we might see that resting-state EEG metrics and cognitive abilities differentially correlate with performance in learning or testing situations. For example, in second language learning, aperiodic activity and IAF has been found to predict neural activity across learning and testing alongside behavioural outcomes (Cross et al., 2020), suggesting information may be differentially integrated and managed with variations in IAF or 1/*f* slope. Further, the 1/*f* slope can indicate an individual’s propensity to improve from practice sessions in a visuomotor task (Immink et al., 2021) and may modulate performance capabilities during learning. As the TMA task is multifaceted and complex, participants engaged in a practice session (to learn the role of the TMA operator) and a testing session of equal length, allowing us to determine how both neural and cognitive metrics dynamically predict performance over time.

We used linear mixed-effects models with cubic splines to model the (non-linear) relationship between IAF, 1/*f* activity and individual differences in a range of cognitive domains on TMA task performance (in practice and test sessions) over time. We hypothesised that a steeper 1/*f* slope and a higher intercept of the aperiodic component would be associated with overall improved performance, as well as a steeper learning curve relative to those with a shallower 1/*f* slope and lower intercept. As TMA involves integrating rapidly changing visual and auditory information to derive solutions, we also expected that a higher IAF would be related to improved overall performance across practice and testing sessions. Further, we hypothesised that the predictive effects of specific cognitive skills (as measured by traditional cognitive tests, e.g., mental rotation) would depend on how closely each test parallels the demands of the TMA task.

## 2. METHOD

### 2.1. Participants

Forty-four adults (M_age_ = 24.30, range: 18 – 40) volunteered to participate in the current study. All participants reported no history of psychiatric, neurological, cognitive or language disorders, normal to corrected-to-normal vision, right-handedness and no use of medications that may affect EEG. Ethics approval was acquired from the university’s ethics committee (project no. 203001) prior to the study’s commencement. Participants received an AUD$60 research honorarium. Due to technical errors during the TMA simulation (no auditory input and incorrect simulation scenario), two participants were removed from the analysis. A further three participants were excluded due to poor understanding of the task’s requirements (demonstrated by a floor effect in performance metrics). Consequently, a total of thirty-nine participants (15 males, 24 females) were included in the final analyses (M_age_ = 24.34, range: 18 – 40).

### 2.2. Traditional Cognitive Measures

Participants completed three standard computerised cognitive tasks. This included the *Visual Statistical Learning Task* (VSL; Siegelman et al., 2017), *Digit Span Task* (DST; Wechsler, 2008) and a *Mental Rotation Task* (MRT; Shephard & Metzler, 1971) to assess visual statistical learning ability, working memory capacity and spatial imagery proficiency, respectively. The VSL task familiarises participants with a series of 16 black-and-white shapes randomly organised into triplets, followed by 42 questions on recognising and completing the triplets. The DST is a common neuropsychological test from the Wechsler Adult Intelligence Scale (Wechsler, 2008) where participants are presented with a series of digits and asked to recall the sequence in the order they appeared. For this experiment, the DST was adapted to involve visual stimuli using OpenSesame (v.3.2.8; Mathôt et al., 2011). The MRT created for this study included 32 alpha-numeric stimuli (digits and letters), 60 2D stimuli (abstract shapes) and 94 3D (polygons) stimuli presented in E-Prime (version 3.0; Psychology Software Tools, 2016) and was based on the parameters described in Jordan et al. (2001).

### 2.3. Target Motion Analysis Task from the Control Room Use Simulation Environment

The dual-screen TMA task (see Figure 2A) within CRUSE (Michailovs et al., 2021) was the complex and dynamic cognitive task used in the current study. In the simulation, participants were required to integrate multiple sources of sensor information for each of the submarine’s surrounding vessels (contacts) to develop its ‘solution’ (an estimate of its location and movement). The result was a model of all the surface contacts or a ‘tactical picture’.

**Figure 2.**
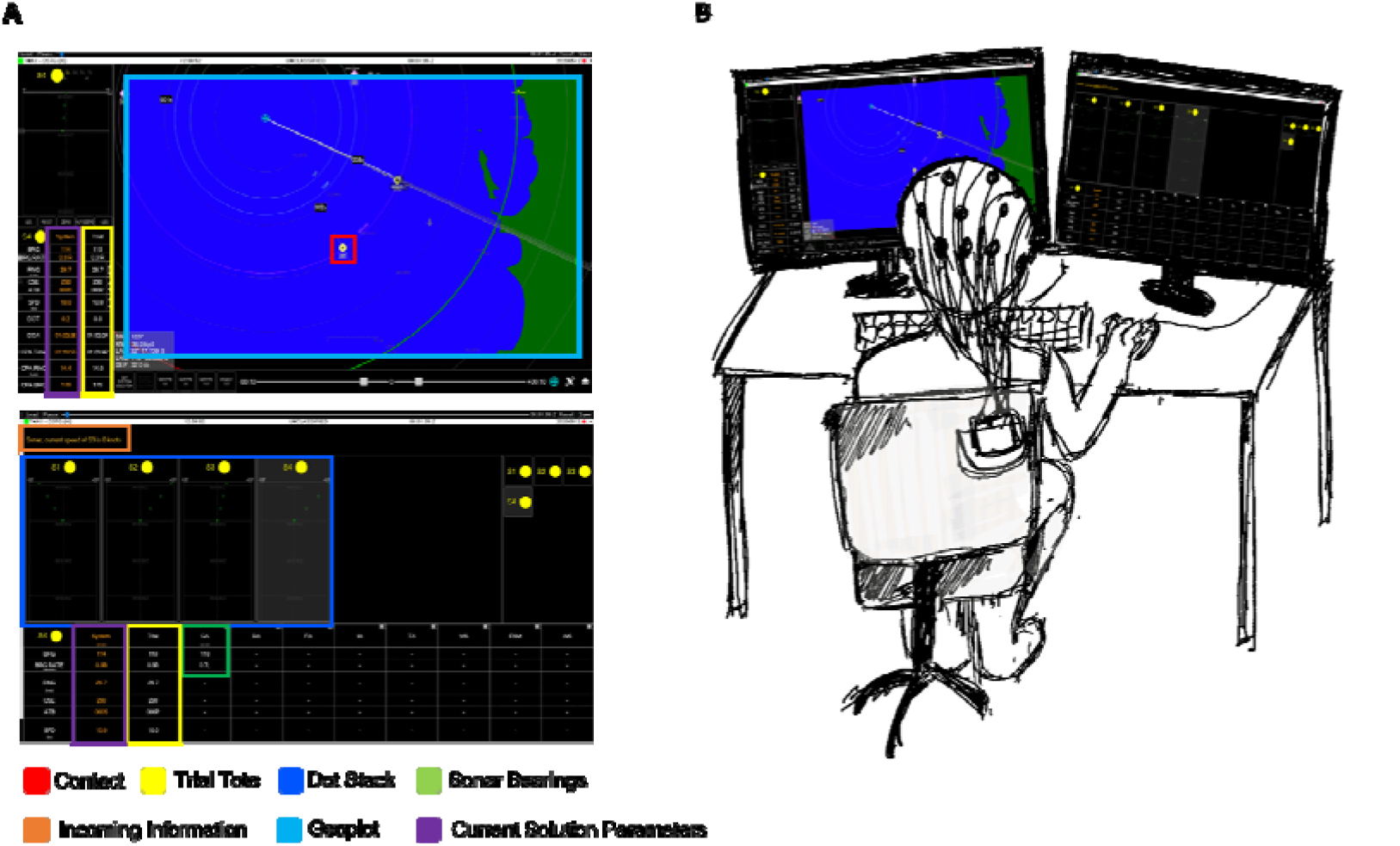
Schematic of the TMA simulation (within CRUSE; Michailovs et al., 2021). **(A)** Left (top) and right (bottom) monitor contents during the simulation. On the left monitor, participants are presented with a geoplot of their submarine, their trial tote (for adjusting contact parameters and setting new solutions) and current plotted solutions. On the right monitor, participants can view the dot stacks and compare their parameters with information received from SONAR, to assess the accuracy of their current solutions. **(B)** Schematic of participants engaging in the dual-screen TMA task within CRUSE.

On the left monitor (see Figure 2A), participants were presented with a geoplot of their submarine’s location and current plotted contact solutions. During the simulation, participants were presented with auditory and visual information representing one or more of: a contact’s position relative to the submarine (bearing), distance from the submarine (range), direction of travel (angle on the bow) or speed. Sensor information would typically be provided by other operators or systems, but was automated in this study. Solutions were derived by adjusting the range, direction and speed in the trial tote and then confirming the solution (see Figure 2B for a schematic of a participant engaging in the TMA task). At any given time, the tactical picture only represented the information entered by the participant, and not necessarily the ‘truth’ of the simulation (i.e., the contact’s objective location as dictated by the simulation’s scenario). Participants could, however, monitor how their solutions were tracking by attending to a panel that visually depicted the solution error relative to information known to be true (i.e., from SONAR) via ‘the dot stacks’ on the right monitor.

We created two TMA scenarios: (1) a practice session with two contacts and standard contact movement, and (2) a testing session, with a markedly higher difficulty, containing five contacts and erratic contact movement. Both the practice and test scenarios were standard across participants. Individual performance was estimated every twenty seconds in the form of tactical picture error (TPE). TPE is a weighted average of all contact position errors (i.e. the distance between coordinates of each solution and the simulation’s ‘truth’ of the surrounding environment) at any point in time (with lower TPE reflecting better performance). Contact position errors were weighted by the contact’s relative priority, a value based on the classification (i.e. warship, fishing vessel), range, course and behaviour (direct course or moving erratically; see Michailovs et al., 2021). The scenarios’ default solutions were included in the calculation of TPE (with all participants commencing the task with a TPE value of 17,500).

### 2.4. Protocol

Interested individuals contacted the researchers to discuss their eligibility. After completing initial screening via email, eligible participants were given access to a TMA education video and a brief quiz on the content, to ensure the task was understood. When the pre-experiment questionnaires were completed (around 50 minutes), participants were invited to the Cognitive Neuroscience Laboratory at the University of South Australia’s Magill campus for a 3.5-hour testing session. On the day of their visit, participants provided written consent to participate and following EEG cap fitting, completed two minutes of resting state EEG recording with eyes closed. Each participant was then randomly allocated to the TMA task or cognitive testing session first.

In the cognitive testing session, participants completed the DST, VSL task and MRT in a randomised order (taking approximately 50 minutes). In the TMA session, participants first engaged in a 40-minute practice scenario to become familiar with the simulator, where the researcher provided step-by-step instructions and feedback on the participant’s role as the TMA operator. These instructions were standardised for all participants (for the practice session script, please see the supplementary material; S1). Following the practice session, the researcher interviewed the participant on their knowledge of how to perform TMA in CRUSE without providing feedback. Participants then engaged in the 40-minute testing session, where the researcher no longer provided feedback. Following the completion of both the TMA task and cognitive testing, participants were provided with an honorarium. Refer to Figure 3 for diagram of the in-laboratory testing session.

**Figure 3.**
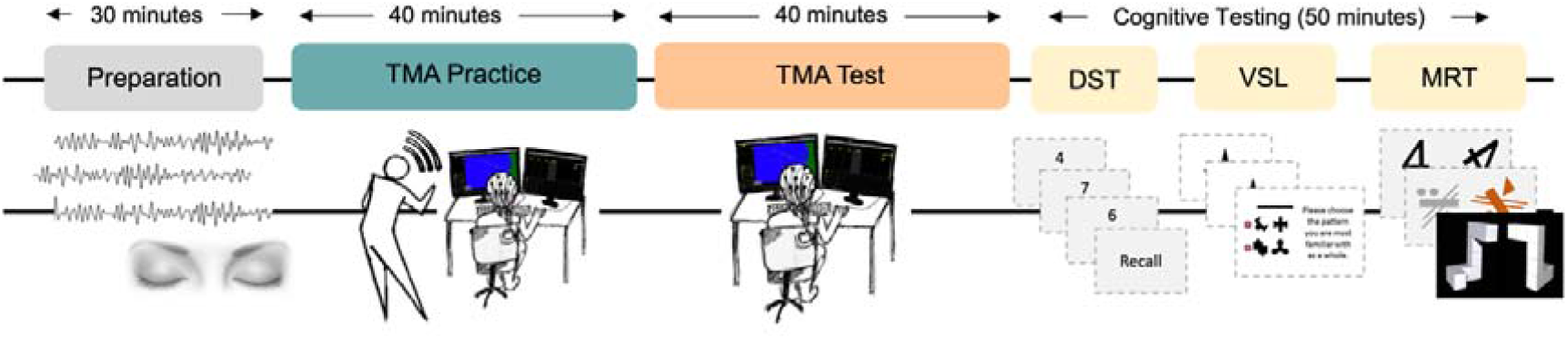
Schematic of in-laboratory testing session protocol. Participants were allocated 3.5 hours to be fitted with an EEG cap, engage in a practice and test session of the CRUSE TMA task and complete the computerised cognitive tasks. Participants first completed a two-minute eyes closed resting-state EEG recording before engaging in either the TMA task (practice and test) or cognitive testing. The order of these two sessions were counterbalanced across participants. In the cognitive testing component, participants also completed the digit span task (DST), visual statistical learning (VSL) task and a mental rotation (MRT) task in a randomised order.

## 3. DATA ANALYSIS

### 3.1. EEG recording and pre-processing

Due to COVID-19, electrophysiological metrics were calculated from EEG recorded at rest using a combination of standard face-to-face EEG collection procedures and archival data. EEG recordings used either a LiveAmp or actiCHamp (Brain Products GmbH, Gilching, Germany) with 32 or 64 active Ag/AgCl electrodes mounted in an elastic cap (actiCap or BrainCap, Brain Products GmbH, Gilching, Germany). Electrode placement followed the 10/20 system. Horizontal and vertical eye movements and blinks were monitored with electrodes placed above and slightly below the left eye or with frontal channels (i.e., Fp1/Fp2) if electrooculographic (EOG) channels were not present. All channels were amplified using a BrainVision amplifier (LiveAmp 32 or actiCHamp base unit 5001, Brain Products, GmbH) at either a 500 Hz or 1000 Hz sampling rate. All EEG pre-processing and analysis was performed using MNE-Python (Gramfort et al., 2013). Raw data were band-pass-filtered from 1 to 40 Hz (zero-phase, hamming windowed finite impulse response). Data were re-referenced to the average of left and right mastoid electrodes or close proximity to mastoids (e.g., TP9/TP10) and resampled to 500 Hz.

### 3.2. Individual alpha frequency estimation

Individual alpha frequency (IAF) estimates were derived from resting-state EEG recordings. IAFs were estimated from six occipital-parietal electrodes (P3/P4/O1/O2/P7/P8) using philistine.mne.savgol_iaf (Alday, 2019) in MNE-Python (see Corcoran et al., 2018; see also Cross, Santamaria, et al., 2020). This IAF estimation routine uses a Savitzky-Golay filter (frame length = 11 frequency bins, polynomial degree = 5) to smooth the power spectral density (PSD). It then searches the first derivative of the smoothed PSD for evidence of peak activity within a defined frequency interval (here, 7 – 13 Hz).

### 3.3. Spectral decomposition and aperiodic (1/*f*) estimation

To separate oscillatory activity from the aperiodic component in resting-state EEG recordings, we used the irregular-resampling auto-spectral analysis method (IRASA v1.0; Wen & Liu, 2016) implemented in the YASA toolbox (Vallat & Jajacy, 2021) in MNE-Python to estimate the 1/*f* power-law exponent characteristic of background spectral activity. IRASA isolates the aperiodic (random fractal) component of neural time series data via a process that involves resampling the signal at multiple non-integer factors *h* and their reciprocals 1/*h*. Because this resampling procedure systematically shifts narrowband peaks away from their original location along the frequency spectrum, averaging the spectral densities of the resampled series attenuates peak components, while preserving the 1/*f* distribution of the fractal component. The (negative) exponent summarising the slope of aperiodic spectral activity is then calculated by fitting a linear regression to the estimated fractal component in log-log space. For a full mathematical description of IRASA, see Wen and Liu (2016).

### 3.4. Statistical analysis

Statistical analyses were conducted using *R* version 3.6.2 (R Core Team, 2020) with packages *tidyverse* v.1.3.0 (Wickham et al., 2019), *car* v.3.0.8 (Fox & Weisberg, 2019), *effects* v.4.1.4 (Fox & Weisberg, 2019), *lme4* version 1.1.23 (Bates et al., 2015) and *splines* version 4.0.2 (Bates et al., 2015). Plots were created in *R* using *ggplot2* v.3.3.0 (Wickham, 2016) and *corrplot* v.0.84 (Wei, 2021). *lmerOut* v.0.5 (Alday, 2018) was used to produce model output tables, while *ggeffects* v.1.0.2 (Lüdecke, 2018) was used to extract modelled effects for visualisation.

#### 3.4.1. Linear mixed effects models

Data were analysed using linear mixed-effects models fit by restricted maximum likelihood (REML) estimates. Each model took the following general form:

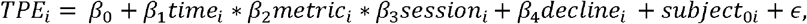

Here, *Time* refers to experimental time in twenty second epochs, *metric* encodes the cognitive (e.g., VSL ability) or resting-state neural (e.g., IAF) metric, and *session* is Practice or Test sessions. *Decline* refers to the regression slope between Time and TPE for the first 500 seconds of the Practice and Test sessions to control for individual differences in performance at the beginning of each task. This parameter was extracted given that participants’ decline in TPE estimates was highly similar in the first portions (500 seconds) of the Practice and Test sessions, as shown in Figure 4A. In order to reduce autocorrelation of the residuals driven by the highly restricted range of responses, this parameter was entered into all models as a covariate. Subject was also modelled as a random effect on the intercept, while TPE was specified as the outcome. Asterisks denote interaction terms, including all subordinate main effects; pluses denote additive terms.

**Figure 4.**
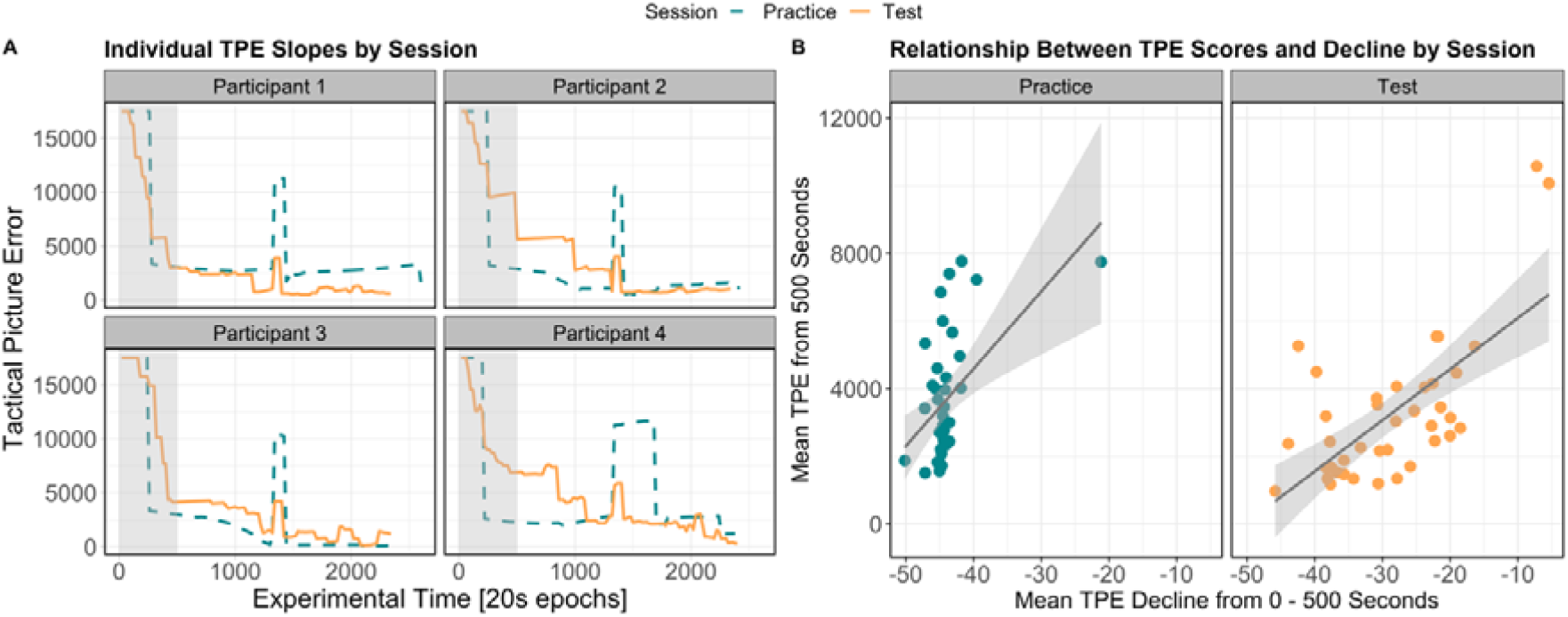
Visualisation of TPE scores in the TMA task across both Practice and Test sessions. **(A)** TPE scores (y-axis; higher scores indicate worse performance) across experimental time (x-axis; left to right indicates the start to end of the task) for a subset of participants, illustrating similarities in TPE scores between the first 500ms (indicated by the shaded regions). **(B)** Relationship between TPE decline (y-axis; regression slope of TPE in the first 500ms) and mean TPE scores in the remainder of the session (x-axis). Shaded regions indicate ± the standard error of the mean.

Type II Wald χ^2^-tests from the *car* package (Fox, 2011) were used to provide *p*-value estimates, while categorical factors were sum-to-zero contrast coded, such that factor level estimates were compared to the grand-mean (Schad et al., 2020). Further, an 83% confidence interval (CI) threshold was used given that this approach corresponds to the 5% significance level with non-overlapping estimates (Austin & Hux, 2002; MacGregor-Fors & Payton, 2013). In the visualisation of effects, non-overlapping CIs indicate a significant difference at *p* < .05. Finally, smoothing splines were applied to *Time* in each model, in order to model non-linear fluctuations in TPE across the Practice and Test sessions as a function of individual differences in the cognitive and neural metrics.

## 4. RESULTS

Overall, participants performed better on the Test (M_TPE_ = 4848, SD = 1975, range: 2594 – 11400) than the Practice (M_TPE_ = 5093, SD = 1609, range: 3127 – 9183) TMA task. As illustrated in Figure 5A, while TPE scores were higher in the Practice session (indicating greater divergence from the scenario ground truth), there was greater variability in the Test session when aggregating across the task; however, as is clear from Figure 5B, when examining performance over time, TPE scores decreased at a faster rate and had lower variability in the Test than the Practice session, indicated by the smaller standard error (i.e., shaded area). For a summary of descriptive statistics of performance on the cognitive tasks, see Table 1.

**Figure 5.**
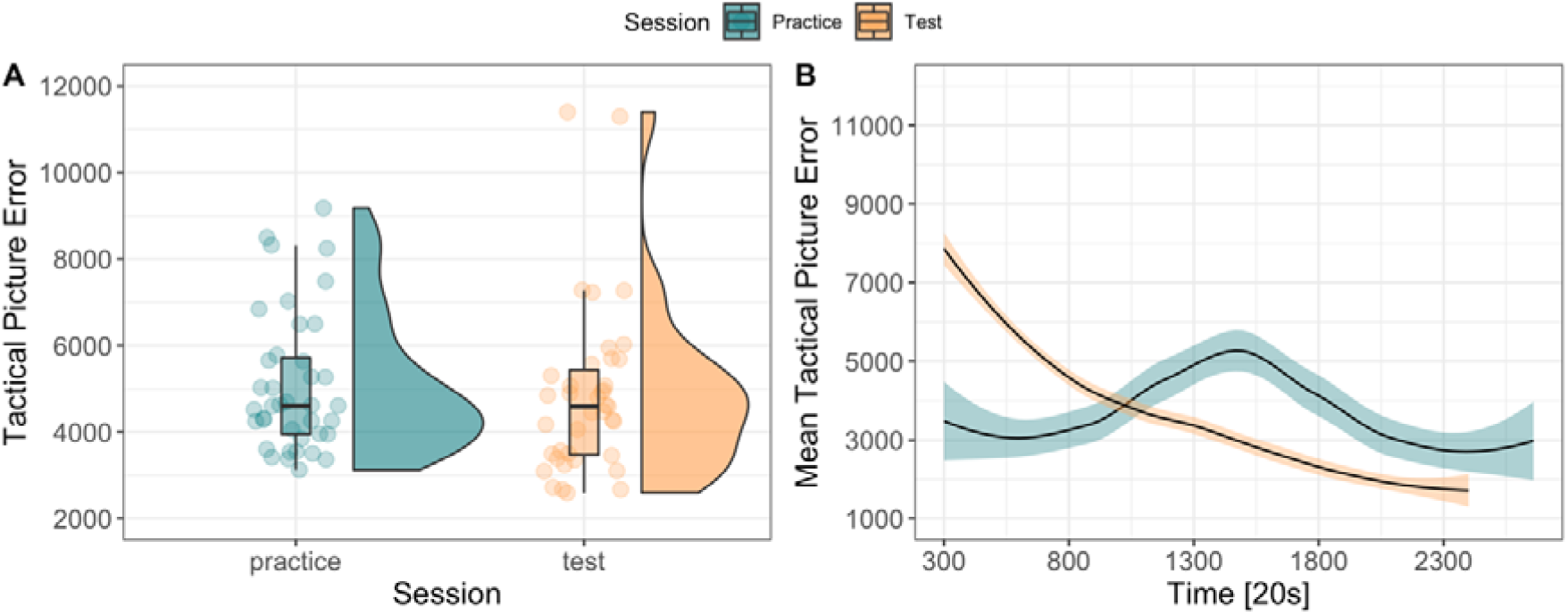
Summary of TPE scores in the TMA task across session and time. **(A)** Raincloud plot (Allen et al., 2019) illustrating average TPE scores between the Practice and Test sessions. **(B)** Average TPE scores (y-axis; higher scores indicate worse performance) across time (x-axis; left to right indicates the start to end of the task, respectively) between the Practice and Test sessions. Shaded regions indicate ± the standard error of the mean.

**Table 1.** Mean, standard deviation (SD) and range in performance for the cognitive tasks.

| Cognitive Task |  |  |  |
| --- | --- | --- | --- |
|  | Mean | SD | Range |
| Statistical Learning | 60.94 | 16.76 | 31 – 100 |
| Digit Span | 6.33 | 1.40 | 4 – 9 |
| Mental Rotation Accuracy | 81.17 | 14.06 | 44.68 – 95.74 |
| Mental Rotation RT | 3626.78 | 1040.35 | 908 – 5664 |

*Note.* RT = reaction time in milliseconds.

### 4.1. Modelling individual differences in cognitive ability to predict performance across time

Here, we examined how performance on the TMA task changes across time between the Practice and Test sessions as a function of individual differences in cognition. The first model focussed on VSL ability, revealing a significant interaction of the cubic time term, session and VSL ability on TPE (χ^2^(4) = 19.2, *p* <.001). As shown in Figure 6A, a higher VSL ability was related to more consistent TPE scores across the Practice, when compared to low and moderate levels of statistical learning ability. However, for those with lower statistical learning ability, performance was greater during most of the Practice session, except for a sudden sharp increase when a second contact was added into the simulation (see Figure 6A, epoch 1500). No modulation by statistical learning ability were observed in the Test session.

**Figure 6.**
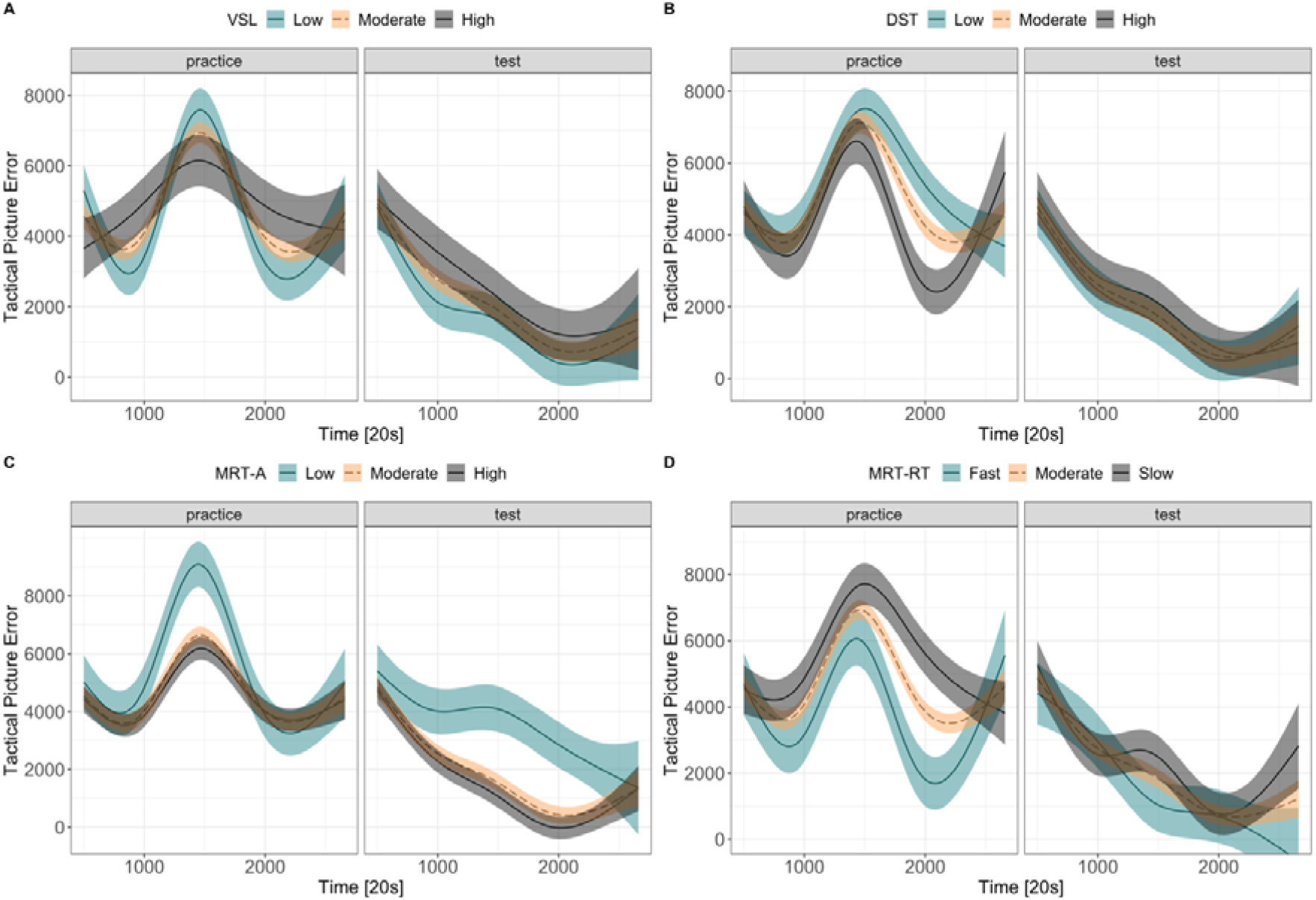
Modelled effects for changes in TPE on the TMA task based on performance on the visual statistical learning (VSL) task (**A**), digit span task (DST) **(B)**, accuracy on the mental rotation task (MRT-A) **(C)** and reaction time on the mental rotation task (MRT-RT) **(D)**. TPE is represented on the y-axis (higher scores indicate worse performance), while experimental time is represented on the x-axis. Facets represent session, with the Practice session on the left and the Test session on the right. Differences in performance on the cognitive tasks are categorised as low, medium, and high or fast, moderate, and slow based on the first quartile, median and third quartile, respectively. Statistical models included these predictors as continuous variables, however, categories were created for visualisation purposes.

Next, we examined whether differences in working memory (quantified using the digit span task) predicted modulations in TPE scores across sessions. There was a significant interaction of the cubic spline term of time, session and digit span scores on TPE (χ^2^(4) = 21.95, *p* <.001). During Practice, a high working memory ability was associated with reduced TPE, particularly in the last third of the task. For the Test session, no modulation of TPE by working memory ability was observed.

Modelling of mental rotation ability (quantified as accuracy on the mental rotation task) revealed a significant interaction of the cubic time term, session and mental rotation ability on TPE (χ^2^(4) = 28.05, *p* <.001), whereby a low mental rotation ability was associated with increased TPE throughout the majority of the Practice and Test session, while a high mental rotation ability was associated with reduced TPE. By contrast, modelling of mental rotation processing speed (quantified as reaction time on the mental rotation task) revealed a significant interaction of the cubic time term, session and mental rotation reaction time (χ^2^(4) = 25.06, *p* <.001). Here, a slower relative to moderate and fast reaction time on the mental rotation task predicted an increase in TPE for the majority of the Practice session. For full model summaries, see the supplementary material (S2).

Taken together, individual differences in cognitive abilities modulate performance over time on a complex cognitive task: a high mental rotation ability was associated with a steeper performance curve (i.e., a faster reduction of TPE) under more complex testing scenarios. Working memory capacity showed differential effects in the Practice and Test session, possibly reflecting the changing demands that learning versus executing the TMA task places on memory. Next, we examine whether individual differences in resting-state oscillatory and aperiodic activity modulate performance over time and between sessions on the TMA task.

### 4.2. Modelling individual differences in resting-state neural activity to predict performance across time

Here, we modelled individual differences in resting-state-derived aperiodic (1/*f* slope and intercept) and oscillatory (IAF) activity to determine whether inter-individual differences in intrinsic neural activity predict performance on a complex cognitive task. For a summary of the raw EEG metrics and for the modelled effects, see Figures 7 and 8. For a summary of the relationship between cognitive and neural metrics, see Figure 8.

**Figure 7.**
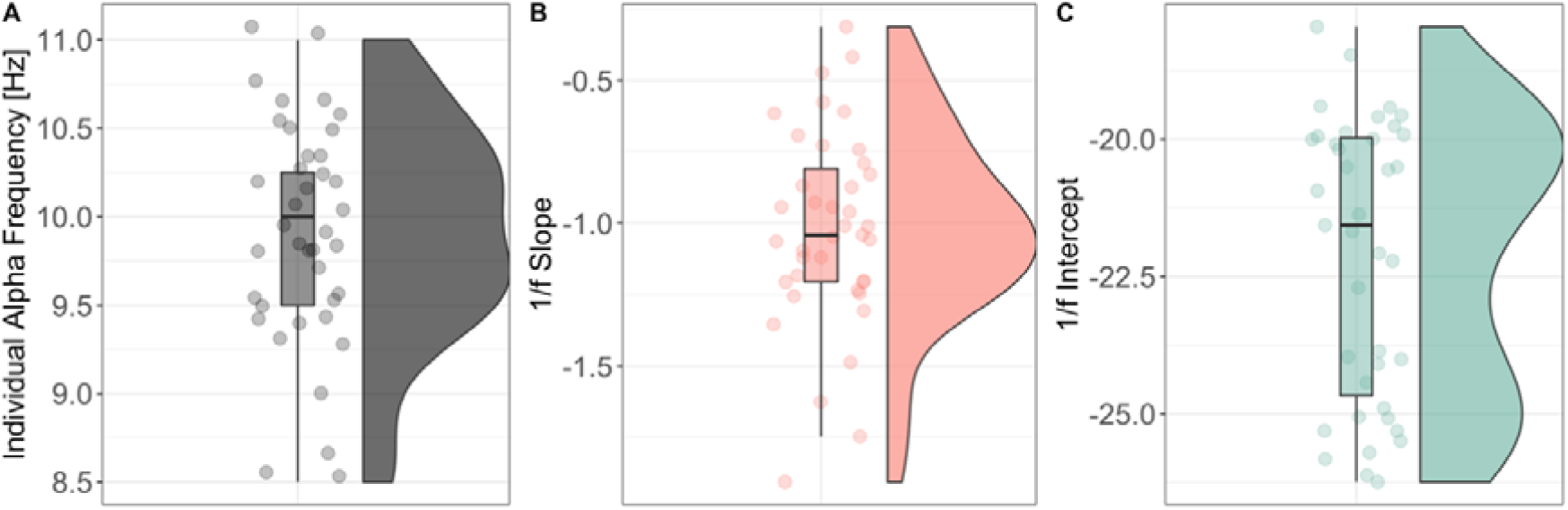
Summary of **(A)** individual alpha frequency (IAF), **(B)** 1/*f* slope and the **(C)** 1/*f* intercept estimated from two minutes of resting-state EEG. Data points indicate individual subject estimates. Thick horizontal lines indicate the median; lower and upper hinges correspond to the first and third quartiles, respectively; lower and upper whiskers extend to the furthest estimate within 1.5 × interquartile range from the lower and upper hinges, respectively.

**Figure 8.**
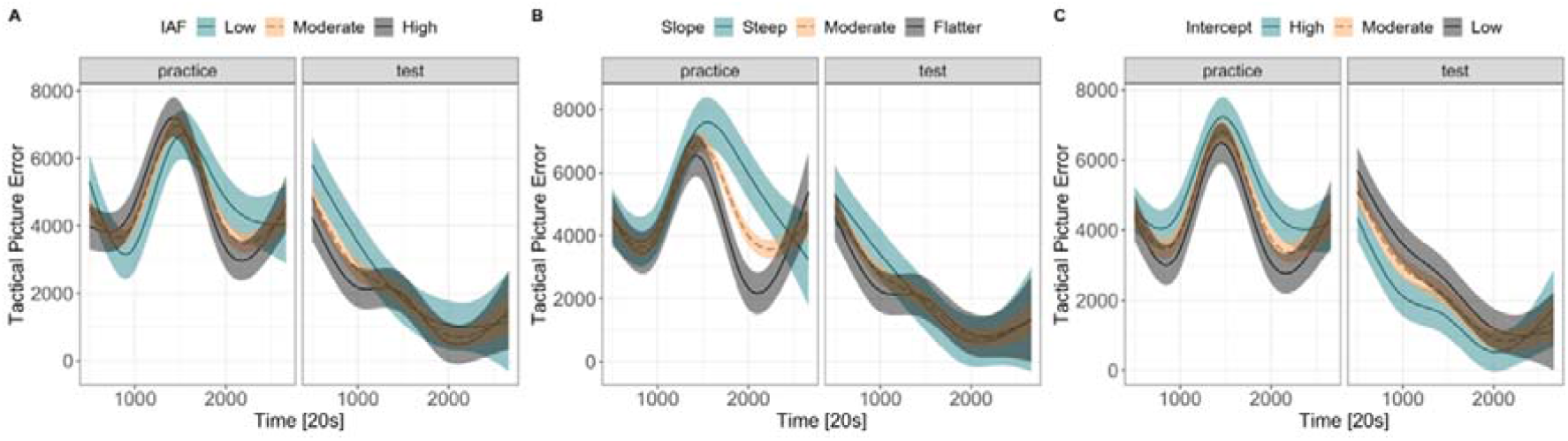
Modelled effects for changes in TPE on the TMA task based on resting-state-derived IAF (**A**), aperiodic 1/*f* slope **(B)**, and aperiodic 1/*f* intercept **(C)**. TPE is represented on the y-axis (higher scores indicate worse performance), while experimental time is represented on the x-axis. Session is facetted with Practice on the left and Test on the right. Differences in resting-state EEG metrics are categorised as low, medium, and high or steep, moderate, and shallow based on the first quartile, median and third quartile, respectively. Statistical models included these predictors as continuous variables, however, categories were created for visualisation purposes.

**Figure 9.**
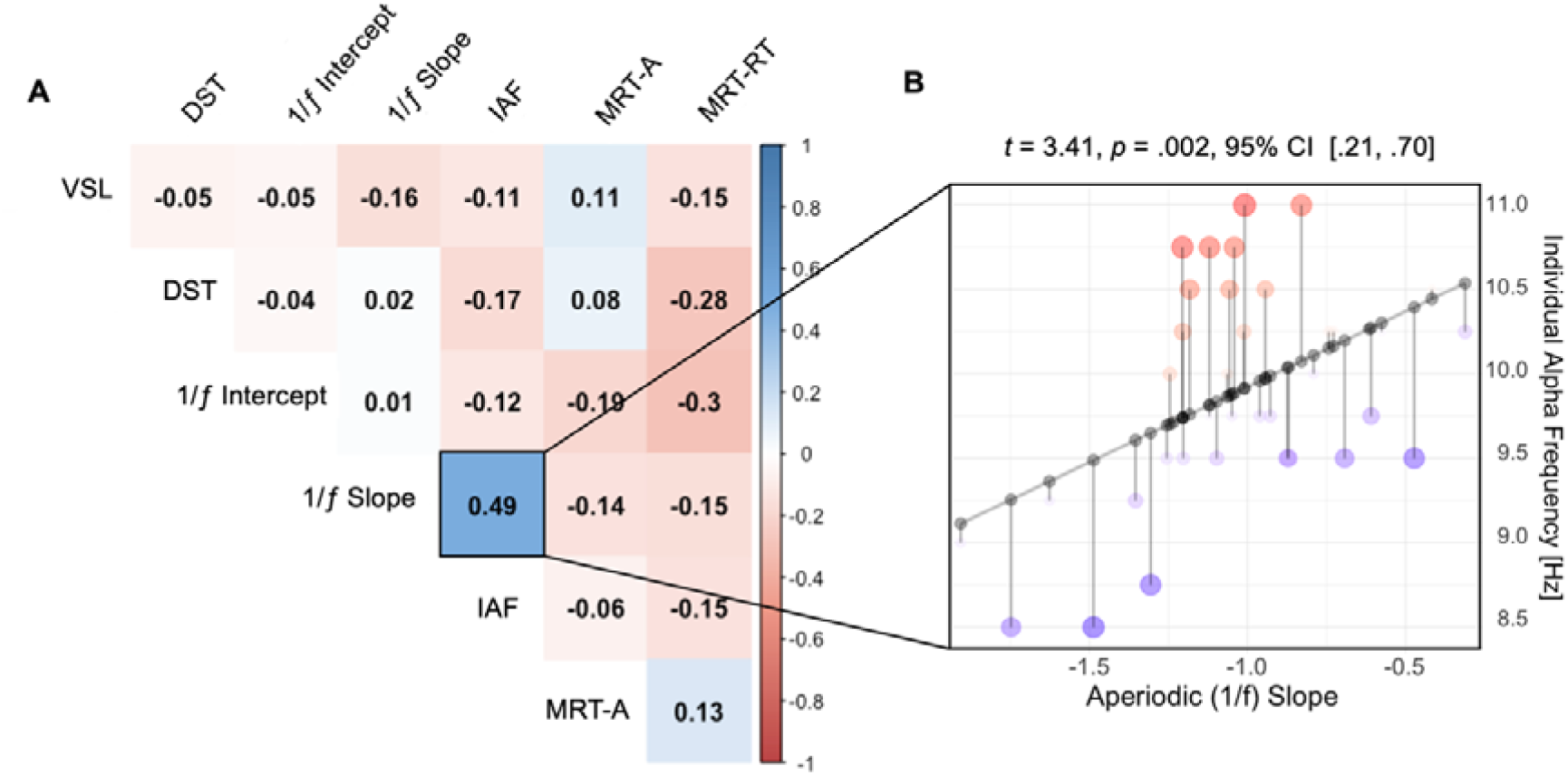
Relationship between cognitive and neural metrics. **(A)** Correlogram illustrating the correlation coefficients between each of the cognitive and resting-state-derived neural metrics. Negative (warmer) values indicate a negative association, while positive (cooler) values indicate a positive association. Values denote person’s *r* coefficients. MRT-RT = mental rotation task reaction time; VSL = visual statistical learning ability; MRT-A = mental rotation task accuracy; IAF = individual alpha frequency. Note that the only statistically significant association was between IAF and the 1/*f* slope. **(B)** Scatterplot illustrating the significant positive association between IAF (y-axis; higher values denote a higher IAF) and the 1/*f* slope (x-axis; higher values denote a shallower slope). Points along the line of best fit indicate predicted values, while points off the line represent residual data, with larger points indicating further distance from predicted values.

The analysis of IAF revealed individual differences in IAF differentially modulated TPE across Practice and Test sessions (with a significant interaction of the cubic time term, IAF and session, [χ^2^(4) = 35.75, *p* <.001]). Lower IAF was associated with decreased TPE in the first portion of the Practice session relative to higher IAF estimates. By contrast, higher IAF was associated with decreased TPE in the first portion of the Test session. No modulations by IAF were observed in the latter portions of the Test session. Modelling of the 1/*f* slope also revealed a significant influence of the 1/*f* slope on TPE (with a significant interaction of the cubic time term, slope and session; χ^2^(4) = 50.22, *p* <.001), with a steeper 1/*f* slope predicting increased TPE in latter portions of the Practice session relative to a moderate and shallow slope. For the Test session, a modulation of TPE across time as a function of the 1/*f* slope was not observed. Finally, for the analysis of the 1/*f* intercept, a significant interaction of the intercept and session (χ^2^(4) = 125.01, *p* <.001) revealed that TPE scores were differentially affected by 1/*f* intercept in the Practice and Test sessions. In the Practice session, a lower 1/*f* intercept was associated with decreased TPE scores but a higher 1/*f* intercept was related to lower TPE scores in the Test session. Further, a significant interaction of the cubic time term and 1/*f* intercept (χ^2^(4) = 1029.94, *p* <.05) indicated that while TPE scores were similar across time (when disregarding session), divergences occurred between higher and lower intercepts around the 1000^th^ and 2000^th^ epochs. For full model summaries, see the supplementary material (S2).

Taken together, these results demonstrate that differences in resting-state neural activity are predictive of performance on a complex cognitive task: a higher IAF is associated with superior adjustment to task demands, while a shallower 1/*f* slope is associated with improved performance during practice but not testing conditions. The intercept of the 1/*f* component also showed an association with the TMA task, with enhanced performance in the practice session associated with a lower 1/*f* intercept and greater performance during the testing session related to a higher 1/*f* intercept.

## 5. DISCUSSION

Previous research has identified how resting-state EEG metrics – both oscillatory and aperiodic in nature – relate to performance on cognitive tasks. The current study aimed to extend previous findings in this area by examining resting-state EEG measures in relation to performance on a multifaceted task where participants were required to rapidly integrate and prioritise multi-modal information to make sense of their surroundings. We used linear mixed-effects models with cubic splines to show that individual resting-state metrics (1/*f* and IAF) and specific cognitive abilities (visual statistical learning ability, digit span and spatial imagery ability) differentially predicted performance on a TMA task over time. Most notably, our results showed that a higher IAF and higher 1/*f* intercept were related to superior adjustment (i.e., faster decline in TPE) during the test session of the TMA task. Contrary to our predictions, the 1/*f* slope did not predict TMA performance in the test session, but shallower slopes were related to improved scores in the latter portion of the practice session. Furthermore, we observed differences in how these distinct factors predicted performance during learning (in the practice session) versus the testing session.

### 5.1 Individual differences in discrete cognitive abilities

We used scores from a range of cognitive tests to determine how individual ability in separable cognitive functions (spatial imagery, visual statistical learning ability and working memory) predicted performance in a more dynamic context. Our results showed that a superior visual statistical learning (VSL) ability was associated with more consistent, but overall decreased performance in the TMA task practice session, however, performance in the test session over time showed general improvements for all participants. VSL ability refers to an individual’s proficiency in extracting underlying regularities and patterns in the environment (Siegelman et al., 2017). As the ability to grasp language relies on understanding grammatical regularities, VSL ability most notably exhibits strong relationships with a range of linguistic functions including sentence comprehension (Misyak & Christiansen, 2012), speech perception abilities (Conway et al., 2010) and second language acquisition (Cross, Zou-Williams et al., 2020; Frost et al., 2013). Performance on the TMA task relies upon being able to flexibly adjust to novel incoming information (e.g., regarding contact movement) that may be erratic, and unlikely to follow statistical regularities. Therefore, a higher VSL ability may impair the integration of more nuanced information due to incorrectly imposing rule-based information in contexts that are not strictly rule-governed (for more detailed discussion, see Bogearts et al. 2022). This proposal is supported by our findings that those with a higher VSL ability exhibited greater overall error in their plotted solutions, when compared to those with low to moderate scores on the VSL task. However, while those with poorer VSL ability had lower overall TPE scores, we also observed a sudden decline in performance when complexity of the practice session increased (i.e., a second contact was added). This could perhaps highlight a trade-off, where those with high VSL ability can more readily integrate/attend to regularities in the environment (albeit possibly impeding peak TMA performance). By contrast, individuals with lower VSL ability, while initially disadvantaged when extrapolating behaviours into novel scenarios, later exhibit superior performance in unpredictable contexts.

Secondly, we found that working memory (WM) capacity differentially predicted performance in the practice and test sessions of TMA. WM refers to an individual’s ability to retain and manipulate information for successful task execution (Chai et al., 2018). In the second portion of the practice session (where participants are learning the TMA role and monitoring two contacts), high WM capacity was associated with better performance when compared to lower WM capacity. This distinction between high and low WM capacity in the practice session may relate to differences in the ability to hold and apply new information (Cowan, 2014). For example, while those with higher WM capacity were able to generalise and apply newly learned information about how to perform the TMA task over time, those with low WM capacity appeared to be disadvantaged when becoming familiar with the TMA role during the practice session. When a secondary contact was added to the simulation in the middle of the session, those with higher WM capacity appeared to be able to adjust rapidly and apply newly learned knowledge, while this appeared to take longer for those with lower WM capacity. Further, as exposure time increased, those with low WM capacity may have required reiteration of the task instructions to perform well. However, this effect dissipated in the test session, perhaps highlighting that WM may play less of a critical role in facilitating optimal performance once the requirements of the TMA task have been understood.

Our results also showed that a greater spatial imagery ability – as measured through accuracy on the mental rotation task – was associated with greater overall performance in the TMA practice and test sessions, when compared to moderate and low spatial imagery ability. Spatial imagery ability refers to the ability to represent the relations amongst objects in space, movements of objects and object parts and other complex spatial transformations (Heuer et al., 1986). From this perspective, the connection between the demands of the TMA task and spatial imagery ability are relatively analogous: TMA requires monitoring the position of objects in space, understanding how certain parameters will translate to changes in contact movement and using visual cues to alter a contact’s expected trajectories or speed. Spatial imagery ability is also known to generalise well across closely related tasks (Uttal et al., 2013). Consistent with this, spatial skills are thought to play a key role in achievement and attainment within science, technology, engineering, and mathematic fields (Shea et al., 2001; Wai et al., 2009), directly corresponding to the role that participants must adapt to as the TMA operator.

Overall, these results highlight how higher skill level across discrete cognitive functions does not consistently translate to better performance in more complex and dynamic settings. Even with cognitive skills closely related to the TMA task demands (i.e., spatial imagery), we showed how superior and inferior ability can produce varying performance patterns across learning and testing scenarios.

### 5.2 Individual alpha frequency and differences in the adaptability of information processing

It has been proposed that the individual alpha frequency (IAF) may act as a neural marker of general cognitive ability (Grandy, Werkle-Bergner, Chicherio, Lövdén, et al., 2013; Grandy, Werkle-Bergner, Chicherio, Schmiedek, et al., 2013), with previous literature demonstrating that higher IAF is advantageous to visuo-perceptual sensitivity (Cecere et al., 2015; Samaha & Postle, 2015) and memory performance (e.g., Klimesch, 1999; cf. Cross, Santamaria et al., 2020). Our results are consistent with previous findings, with a higher IAF related to better overall performance in the first portion of the TMA testing session. Here, participants must adapt to increasing task complexity (from monitoring two contacts to four). However, over time, the strength of this relationship weakened, with low, moderate, and high IAF estimates predicting similar performance on the TMA task toward the end of the test session.

This underlying distinction between individuals with high versus low IAF may relate to the inhibitory function of alpha activity. Alpha oscillations are thought to index inhibitory ability and contribute to optimal suppression of task-irrelevant and selection of task-relevant information (Klimesch, 2012). At the commencement of the TMA testing session, better suppression of task-irrelevant information may allow those with higher IAF to adjust to the task demands more quickly (i.e., prioritise one contact at a time). However, while this is initially crucial to performance, participants with a lower IAF may adaptively engage in other preferred strategies to match performance. Further, within the cognitive neuroscience of language literature, there is evidence that IAF reflects individual differences in information processing strategies (Bornkessel et al., 2004; Kurthen et al., 2020), with lower and higher IAF individuals differentially dealing with ambiguous rule-based information and ambiguity resolution. With heightened perceptual sensitivity (Cecere et al., 2015; Immink et al., 2021; Samaha & Postle, 2015) and tendency to update their internal models of the world more often (Kurthen et al., 2020; Surwillo, 1963), those with a higher IAF may adjust more rapidly to novel incoming information, a strategy that is advantagoues to the dynamically changing TMA environment.

From this perspective, the current findings indicate that IAF – as a measure of inter-individual differences in information processing – is predictive of performance on a complex cognitive task, extending previous findings that demonstrate a relationship between IAF and traditional cognitive tests (e.g., IQ measures, attention tasks). As such, our results indicate that higher IAF may be advantageous for faster adaptation to novel conditions in dynamic environments, translating to more rapid optimal performance in complex settings, such as TMA.

### 5.3 Aperiodic activity as a biomarker of inter-individual differences in cognition

While the aperiodic 1/*f* component has largely been ignored or treated as a nuisance variable in neurophysiological research (Donoghue et al., 2020; He, 2014), emerging research is demonstrating its functional relevance in human cognition (Dave et al., 2018; Ouyang et al., 2020; Voytek et al., 2015). In the current study, the 1/*f* slope did not predict performance in the TMA testing session, which is inconsistent with previous literature, where a steeper resting 1/*f* slope is often related to enhanced cognitive performance (Donoghue et al., 2020; Ouyang et al., 2020; Voytek et al., 2015). However, many of these studies correlate resting-state-derived 1/*f* slope estimates with performance on simple cognitive tasks (e.g., processing speed on an object recognition task, Ouyang et al., 2020). Here, we used resting-state 1/*f* slope estimates to predict time evolving performance on a multi-modal and dynamic task. We did, however, observe that a flatter relative to steeper 1/*f* slope was related to improved performance in latter sections of the practice session, a period where participants were required to rapidly learn the role of TMA. A similar pattern of results has been reported in the context of an artificial grammar learning study (Cross, Corcoran, et al., 2020). Here, the 1/*f* slope became flatter over the course of learning, while performance increased as a function of exposure to the language-related rules. Furthermore, Immink et al. (2021) demonstrated that individuals with a shallower slope benefitted most from practice on a complex visuomotor task. From this perspective, while a steeper 1/*f* slope may be related to enhanced performance on simple cognitive tasks, a shallower slope may be beneficial for learning under more dynamic conditions. This interpretation is supported by recent intracranial work (Sheehan et al., 2018), which found that a flatter slope was associated with better performance on a complex associative memory task. A reduction in the steepness of the 1/*f* slope has been suggested to reflect improved adaptability and efficiency of neural systems (Sheehan et al., 2018), and as such, a flatter PSD slope may arise due to the infusion of noise into ongoing neural signals. This is consistent with the observed negative correlation between our participant’s IAF and 1/f slope, whereby higher IAF was related to flatter slopes. Flatter slopes may thus reflect more complex brain activity, which, within the context of TMA, may be critical due to the complexity and richness of the information. From this perspective, more complex brains – indexed by shallower resting-state 1/*f* slopes – may perform better under informationally rich contexts.

Moreover, the 1/*f* intercept showed a clear interaction with performance on the TMA task, with a higher 1/*f* intercept predicting superior performance across a large majority of the TMA test session relative to a lower 1/*f* intercept. The 1/*f* intercept is thought to reflect increased neural population spiking (Manning et al., 2009; Miller et al., 2012), and is positively correlated with the fMRI BOLD signal (Jacob et al., 2021; Winawer et al., 2013). We also recently demonstrated that a higher 1/*f* intercept is predictive of enhanced visuomotor learning (Immink et al., 2021). In this respect, greater overall neural activity across the frequency spectrum, as reflected by a higher 1/*f* intercept, may be beneficial for time evolving performance in TMA. Specifically, greater overall neural activity may be reflective of increased neural network communication under conditions of higher cognitive complexity (Gardony et al., 2017; Jaušovec & Jaušovec, 2000; Mölle et al., 1996), facilitating the flow of task-relevant information within and between task-relevant neural networks in response to dynamic changes in sensory input. This idea accords with early work on EEG complexity during creative thinking (Mölle et al., 1996) and the completion of complex cognitive tasks (Jaušovec & Jaušovec, 2000), showing that increased EEG spectral power tracks increasing task complexity. More recent work using the mental rotation task has also shown that increases in low frequency (i.e., < 7 Hz) power are associated with superior mental rotation ability (Gardony et al., 2017).

### 5.4 Limitations and future directions

While we examined the influence of resting-state derived EEG metrics on TMA performance, which provide insights into inter-individual differences in intrinsic neural activity, we did not examine task-related EEG during the TMA practice and test sessions. Examining task-related modulations in both oscillatory (e.g., alpha activity) as well as aperiodic (e.g., 1/*f* slope) activity may provide insight into more fine-grained changes in neural activity during a complex, multi-dimensional cognitive task. For example, pre-stimulus 1/*f* exponents have been shown to modulate task-related alpha power during a visual spatial discrimination task (Tran et al., 2020) while task-related modulations in the 1/*f* slope and alpha power interact to influence complex rule learning (Cross, Corcoran, et al., 2020). Further, a more targeted analysis of cognitive factors and their interactions with neural indices could help to disentangle the relationships between each of these predictors and how they work in conjunction or in contrast. Additionally, TMA performance may be related to cognitive domains that were not examined here. For example, the cognitive measures we employed were all in the visual modality (e.g., visual statistical learning, mental rotation task); however, TMA may also rely on auditory information (e.g. range of speed estimates provided verbally from sensor operators). From this perspective, the use of an auditory oddball task, for example, may provide insight into inter-individual differences in auditory information processing, with greater oddball detection perhaps being related to more efficient auditory information integration during the TMA task. We also only examined changes TMA performance between the practice and test sessions over a relatively short period. Further extension of this work could examine whether the relationship between TMA performance and resting-state measures are stable over long durations and determine whether there is a relationship between scenario complexity and resting-state metrics in experienced operators (e.g., trained submariners). Moreover, a more fine-grained analysis of the TMA task (i.e., analysis of the time-evolving requirements of the task and subsequent cognitive demands) could help to determine more temporally accurate relationships between performance at a given point in time and relevant cognitive abilities/neural metrics.

### 5.6. Conclusion

The current study examined how resting-state EEG measures (IAF and 1/*f*), along with traditional cognitive measures, predict performance on a dynamic semi-naturalistic cognitive task. Our findings demonstrate that while some closely related cognitive skills were able to predict performance (for example, spatial imagery ability), these effects were distinct across practice and test sessions. Further, we demonstrated that higher IAF is associated with enhanced adjustment to task demands in dynamic settings, suggesting that this resting-state metric may be critical for information processing across a range of tasks and conditions of complexity. Regarding the emerging 1/*f* literature, we did not observe the expected relations between a steeper 1/*f* slope and cognitive performance; however, this difference may be related to individual differences in underlying brain signal complexity, with more complex neural signals, as reflected by a shallower 1/*f* slope, performing better under conditions of increasing complexity. We also demonstrated preliminary evidence that the 1/*f* intercept is related to varying degrees of performance on the CRUSE TMA task, highlighting the functional significance of separable aperiodic components in higher-order cognition.

## Acknowledgements

The authors are indebted to Justin Hill (Royal Australian Navy) for invaluable insights into submarine operations and the TMA role, as well as to David Munro-Ford for adapting CRUSE to meet the needs of our experiment. The authors would further like to thank Dr Samuel Huf for his instrumental role as our liaison with the Defence Science and Technology Group. We are also grateful to all who participated in the study.

