## Supplementary Materials for "Neural and cognitive correlates of performance in dynamic multi-modal settings"

**(S1) CRUSE Practice Session Speech**

As you saw in the video, submarines must have a picture of the environment around them, as they cannot always come up to the surface to view the surrounding vessels (Also known as **contacts** in submariner terms). Developing a tactical picture involves plotting a solution for each surrounding vessel, that is as close to the vessel’s real location as possible. In this task, you’ll be simulating the job of the target motion analyst (TMA) using this program called CRUSE. Remember, the TMA in a submarine must integrate information to predict where the surrounding vessels are. AIM: to use the information given to you from sonar, optics and the track manager in order to estimate the true location of a contact in relation to your submarine. Your solution must be as close to the contact’s true location as possible.

If we look at the screen here, you can see land (green) and water (in blue). This cross is your submarine, which is travelling North (Point to North). These pink circles represent a contact. For example, this one is called ‘Sierra 1.’ A second contact would be called ‘Sierra 2’ and so on. Sonar will typically tell you the classification of the contact once you click on it, but you can also check it by hovering over this yellow circle here (point to dot stack). Each contact in CRUSE has a set course, range, speed, bearing etc (this is its TRUE path, which it will travel on throughout the simulation). However, the only real information that you know is it’s bearing, obtained from sonar here in this panel (View CA panel). For example, a bearing of .... would mean that the contact is in this direction relative to our submarine. As you can see, as the contact moves, a bearing fan begins to form. Each line represents a bearing cut from sonar. For example, at time 1, sonar detected the vessel in this direction, relative to our ship. At time 2, its bearing was detected here and so on. This is the only information that you can be certain of- you still have to estimate the contact’s range, course and speed in order to estimate the true location of the contact

Although you can see the contact here, it will be altered when you change your solution (Alter range and show them that the contact moves). Therefore, this image is a depiction of where you’ve set your solution, not of where the contact actually is.

Now, this may seem like a lot of information but I’ll go through with you **how to actually set a solution** and you’ll have an opportunity to practice. Contact announced, drag and drop. Click contact, zero the solution; this aligns your bearing with the bearing from sonar. If you look here you can see your trial tote, which displays the speed, ATB, course and bearing, which you have to edit in order to change your solution. Once you are happy with your trial solution, you can click ‘set solution,’ in order to update it into the system, and it will come up on the left in orange. That will be the current solution, but you can edit it and set new solutions as much as you like. You might be wondering “How am I supposed to know what these numbers should be?” Well the answer is “you don’t.” Sonar, optics and the track manager will sometimes tell you the speed or range of the contact, so you can start by using that information and entering it into the system.

**Bearing**

Additionally, you should aim to match up your trial bearing with the bearing from sonar as often as possible. As bearing, range, course and speed are all related, changing the other three will enable you to match up the bearing. This will get you close to having an accurate solution However, just because the bearing may be right doesn’t mean that the rest of the information is, so you’ll have to be constantly checking your solution. As you saw in the training video, vessels with the same bearing can have vastly different ranges, speeds and courses. Sonar also tells you the bearing rate here, which is the direction change per minute, and whether the contact is drawing right or left. So for example, if you stood on the top of your submarine and looked at the contact, you could see it drawing left

**Course**

Therefore, this is related to the contact’s direction of travel, altering the COURSE and the ATB here will affect this, enabling you to also match up your bearing rate with that from sonar

**Speed**

You can also see the speed strip, which displays the direction that you’ve decided the contact is moving, and how fast you think they are going. The longer (or more stretched out) the speed strip is, the faster you have deemed the vessel to be going. Therefore, the speed strip is altered whenever you alter your solution (show them what it looks like when you change the speed)

Ideally, you want the speed strip cuts to align with the bearing lines, as this tells you that your solution is tracking well. So as you can see here, ‘R1’ represents ‘time 1,’ so it would make sense to align this with the first bearing cut from sonar. If the ticks and the bearing cuts weren’t aligned, this would suggest that the speed you have deemed the contact to be going does not correspond with the bearing rate that sonar has calculated. However, as the speed strip also shows the course that you have set, if something is rapidly changing direction, it’s okay if your speed strip doesn’t line up with the bearing cuts. You should just use other cues to assess solution accuracy

**Dot Stack**

If we look over at this panel, we can see how well our solution is tracking. The zero line here represents your solution, and the dots represent the true bearing of the contact at certain time intervals. Ideally, you want the dots to line up with the centre line, as this tells you that your solution is tracking well. Each horizontal line here on the dot stack also represents each solution. You can switch through the solutions as often as you like. Clicking on a contact will produce information from sonar and optics, so it’s up to you how often you access it. Once you have more contacts, you should also prioritise how long you spend on each one according to their classification and speed, and how close they are to you. Additionally, remember that not all contacts will travel on a consistent course and speed, for example some fishing vessels can be quite erratic. Finally, even if you set a solution that’s perfect, your submarine is moving so how well you can track the contact will change.

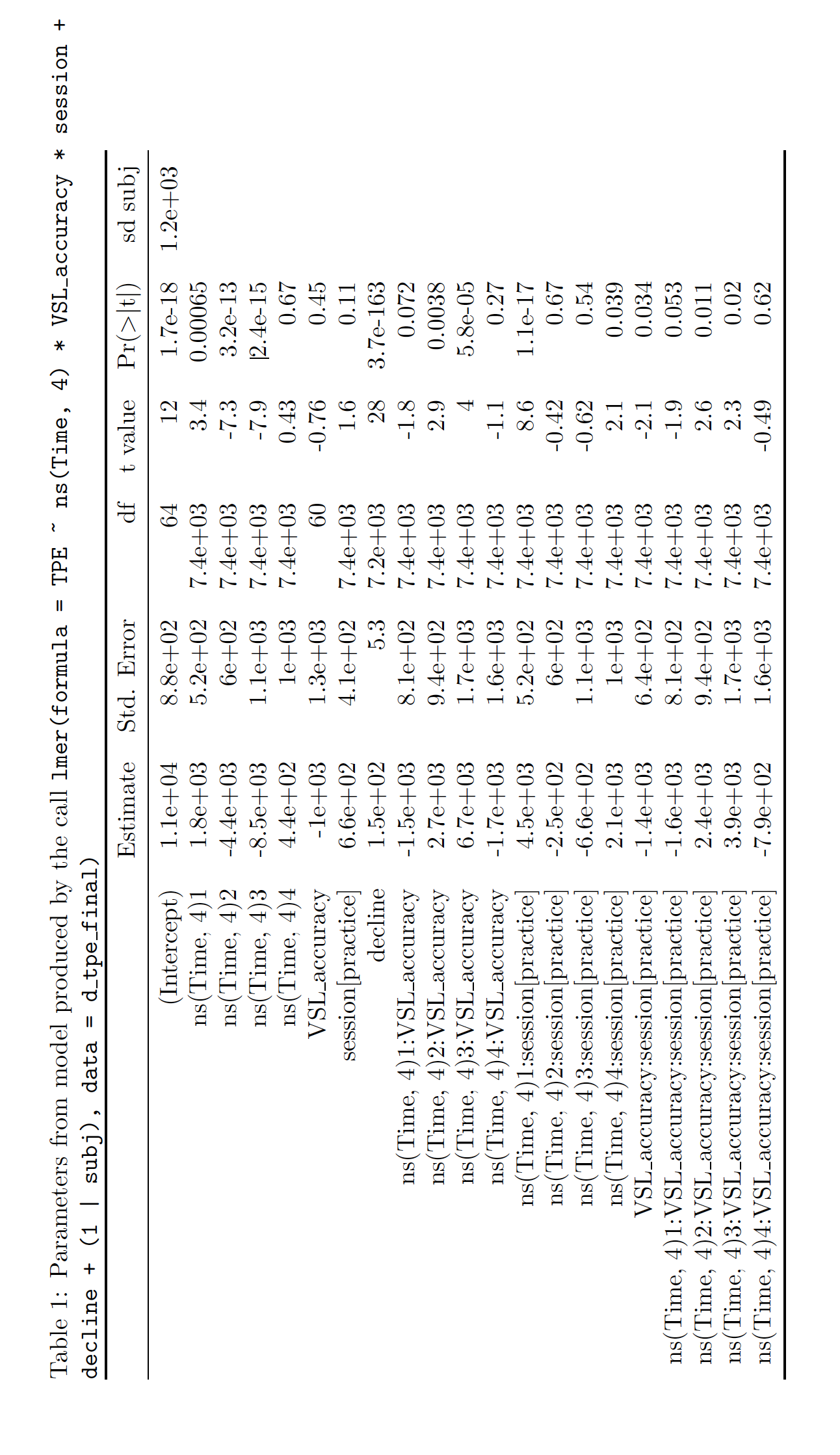
**(S2) Model Outputs**

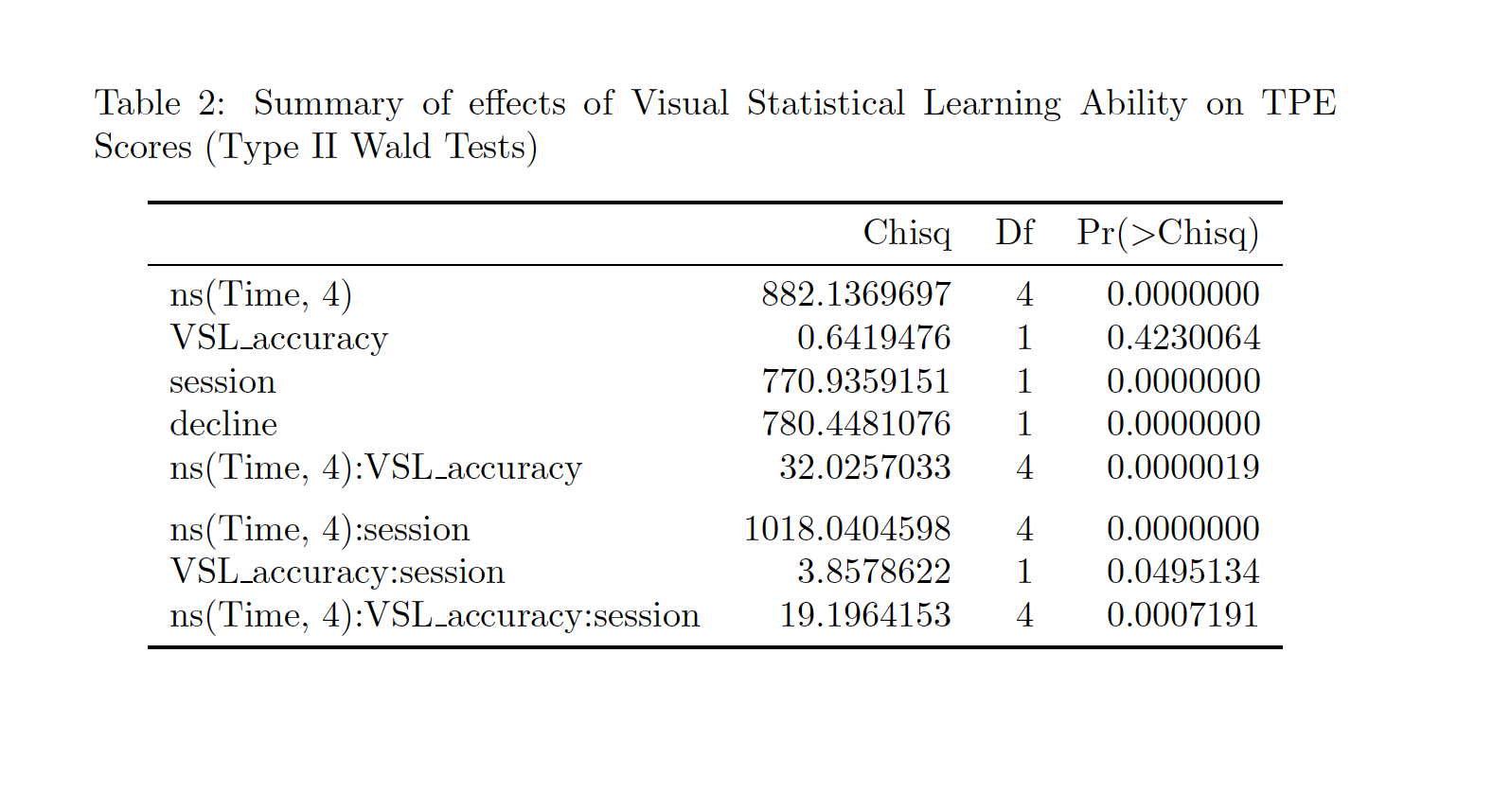

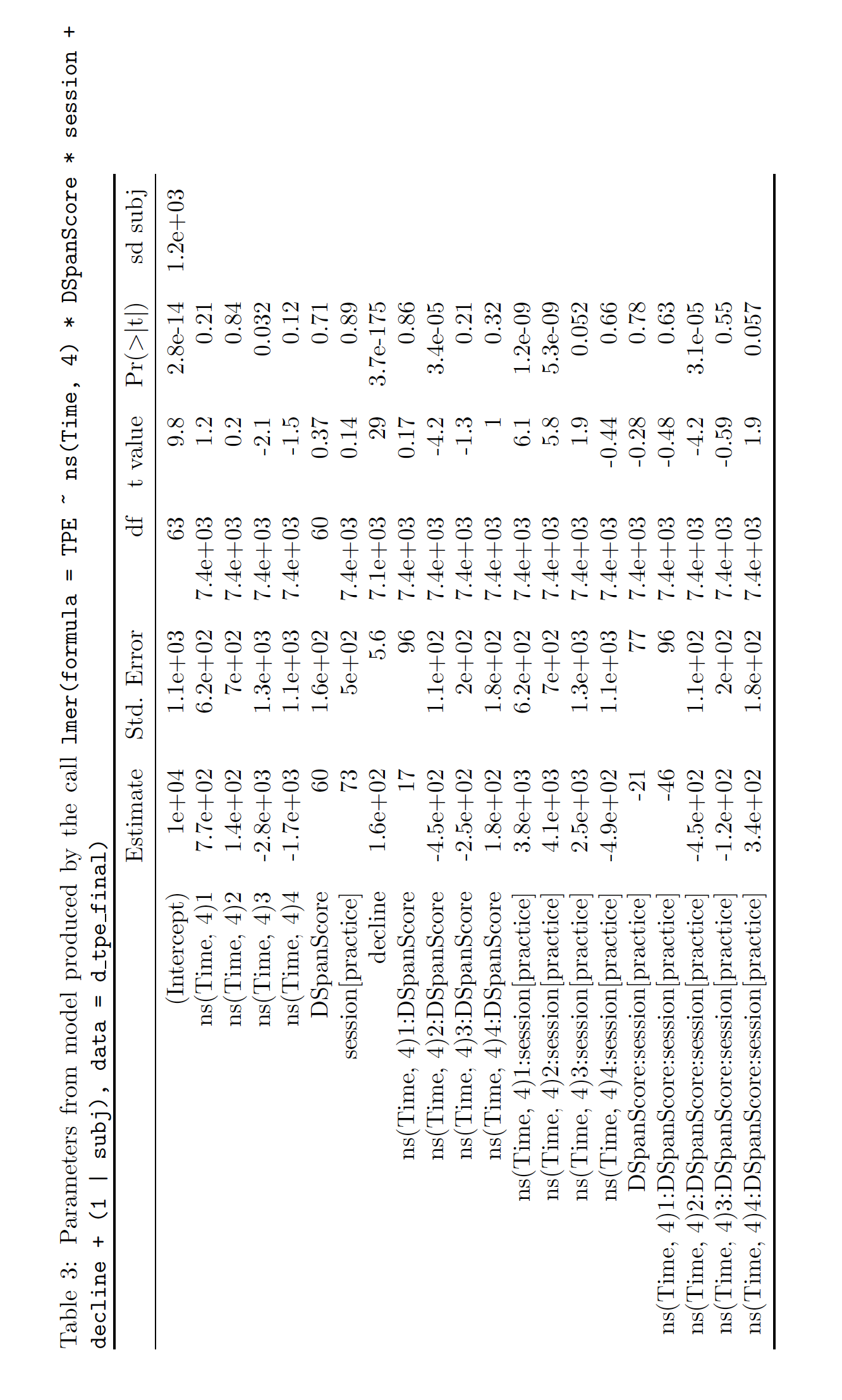

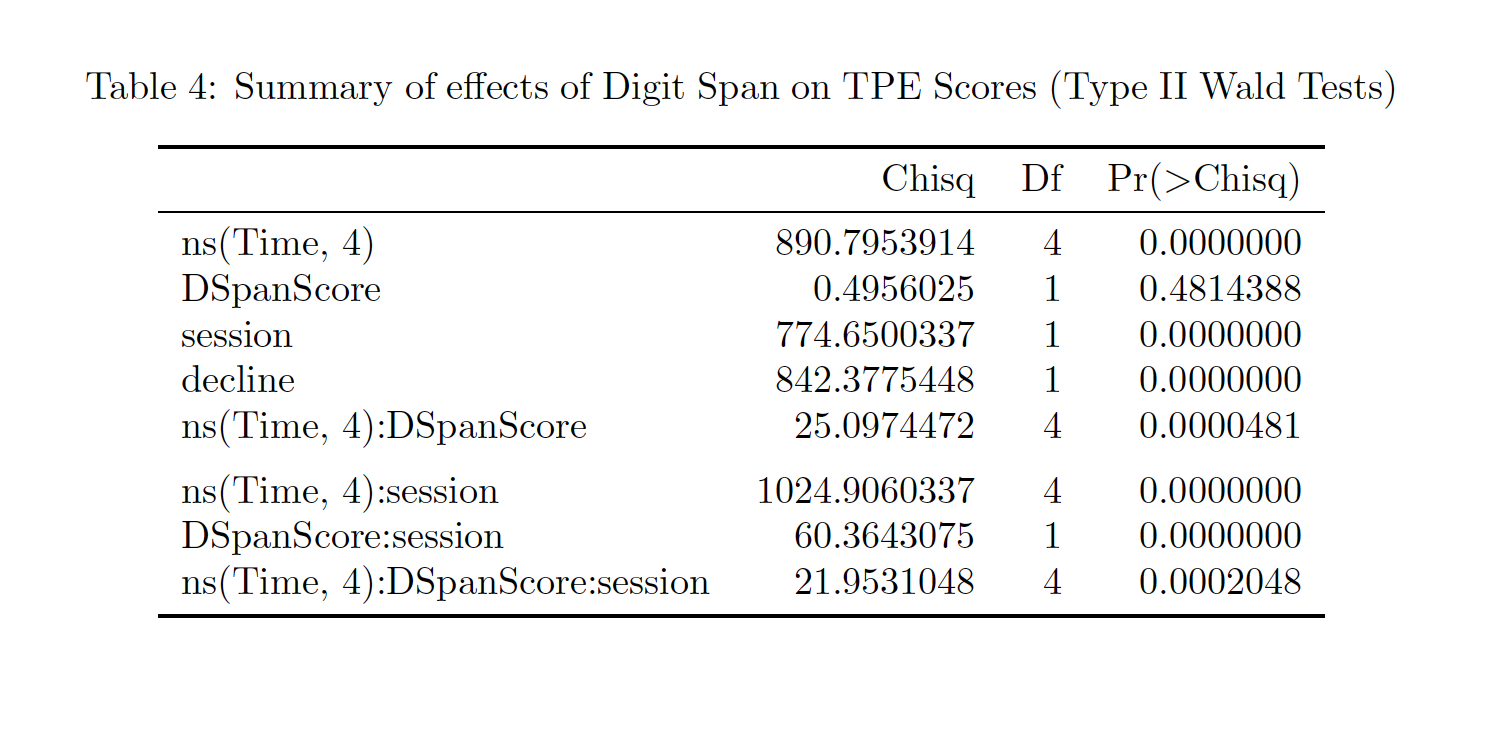

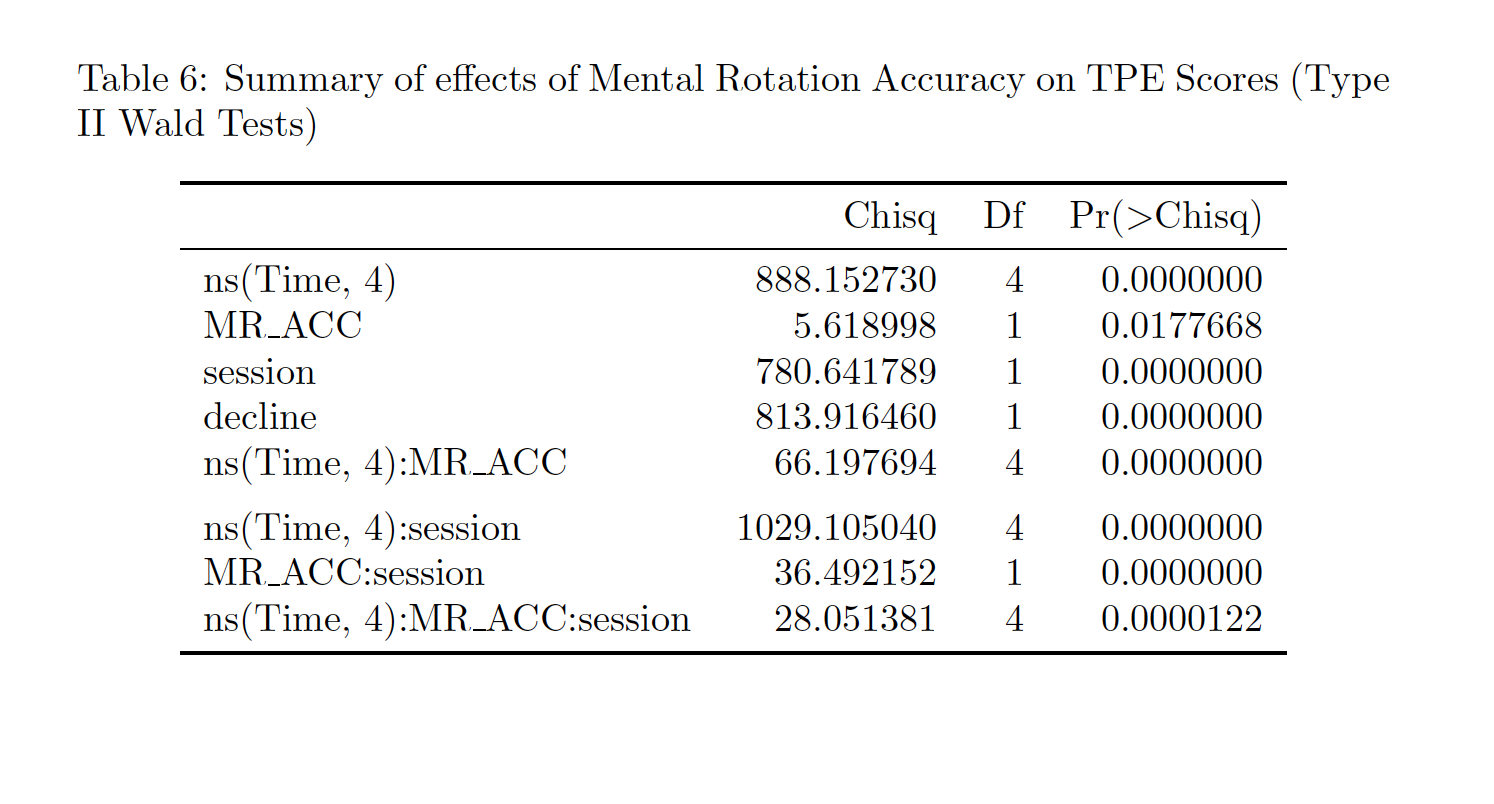

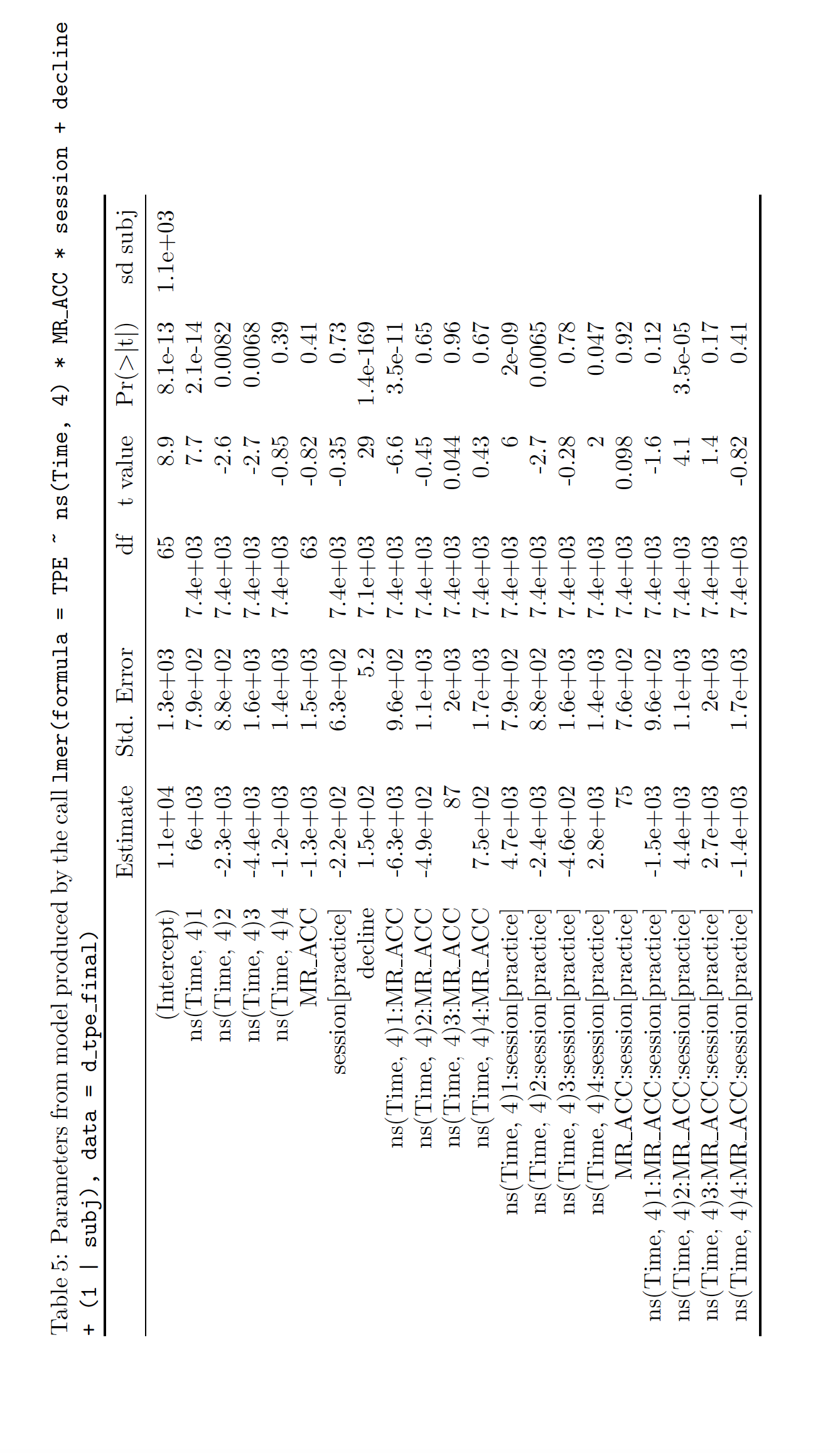

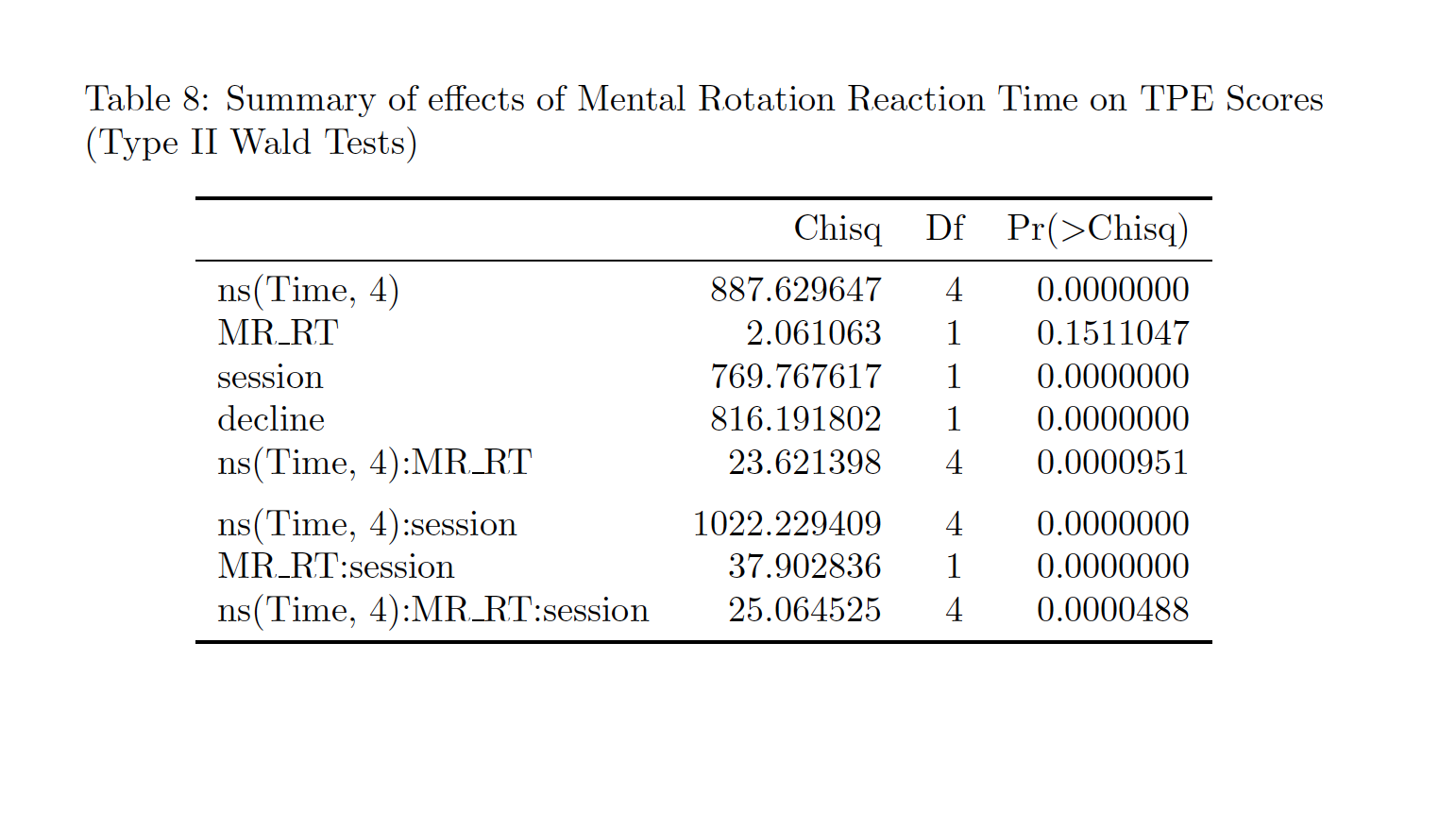

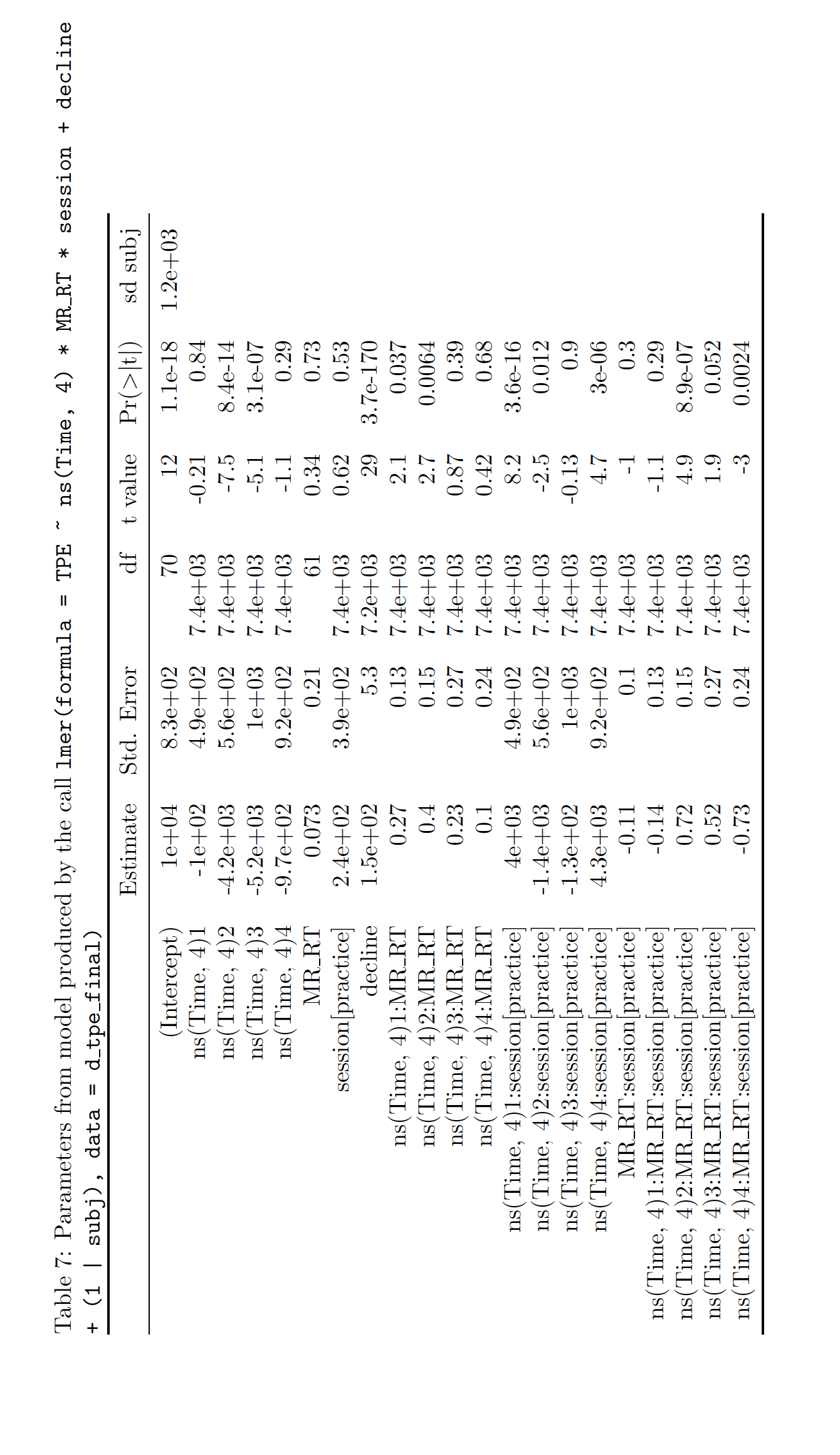

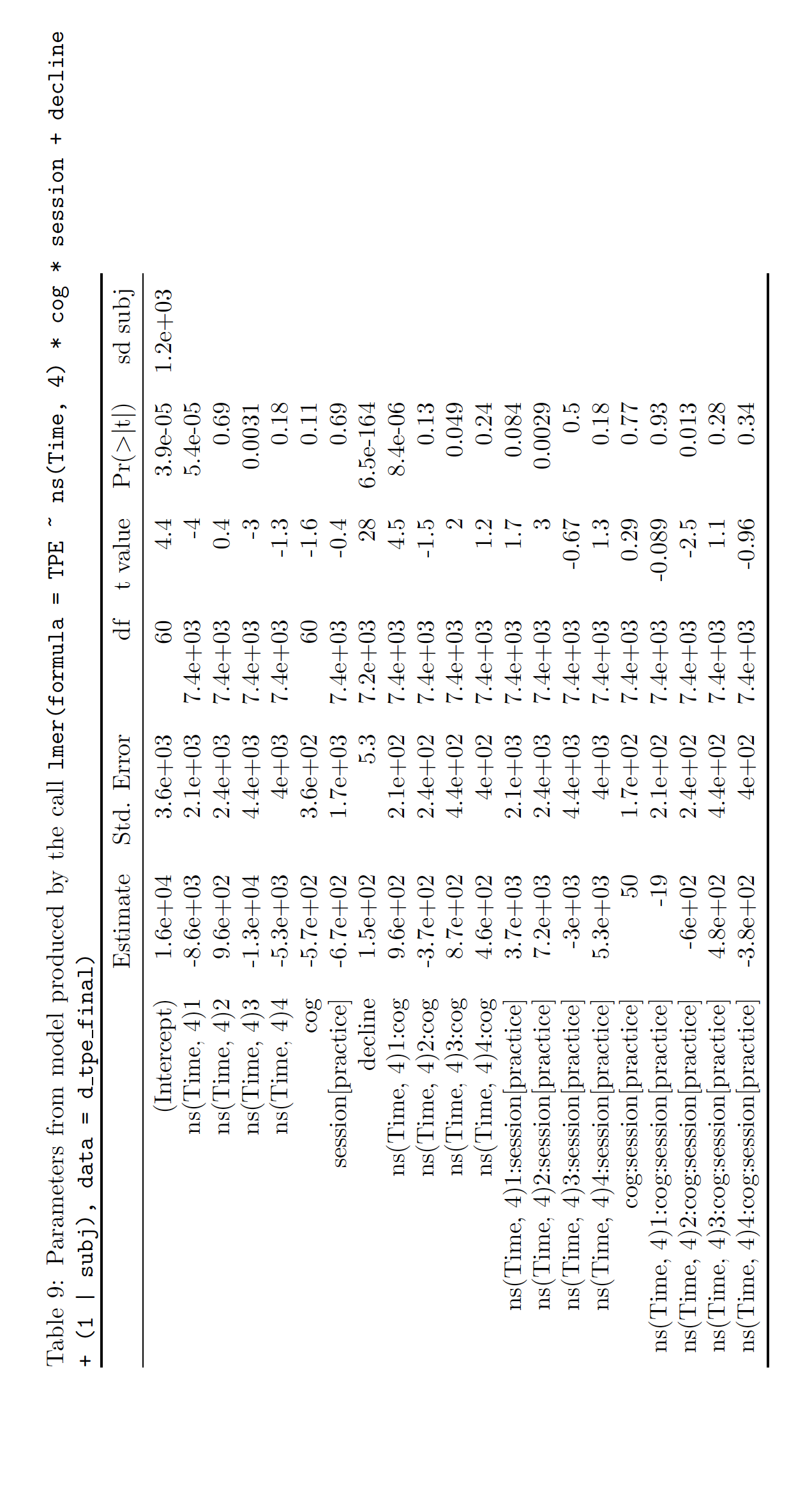

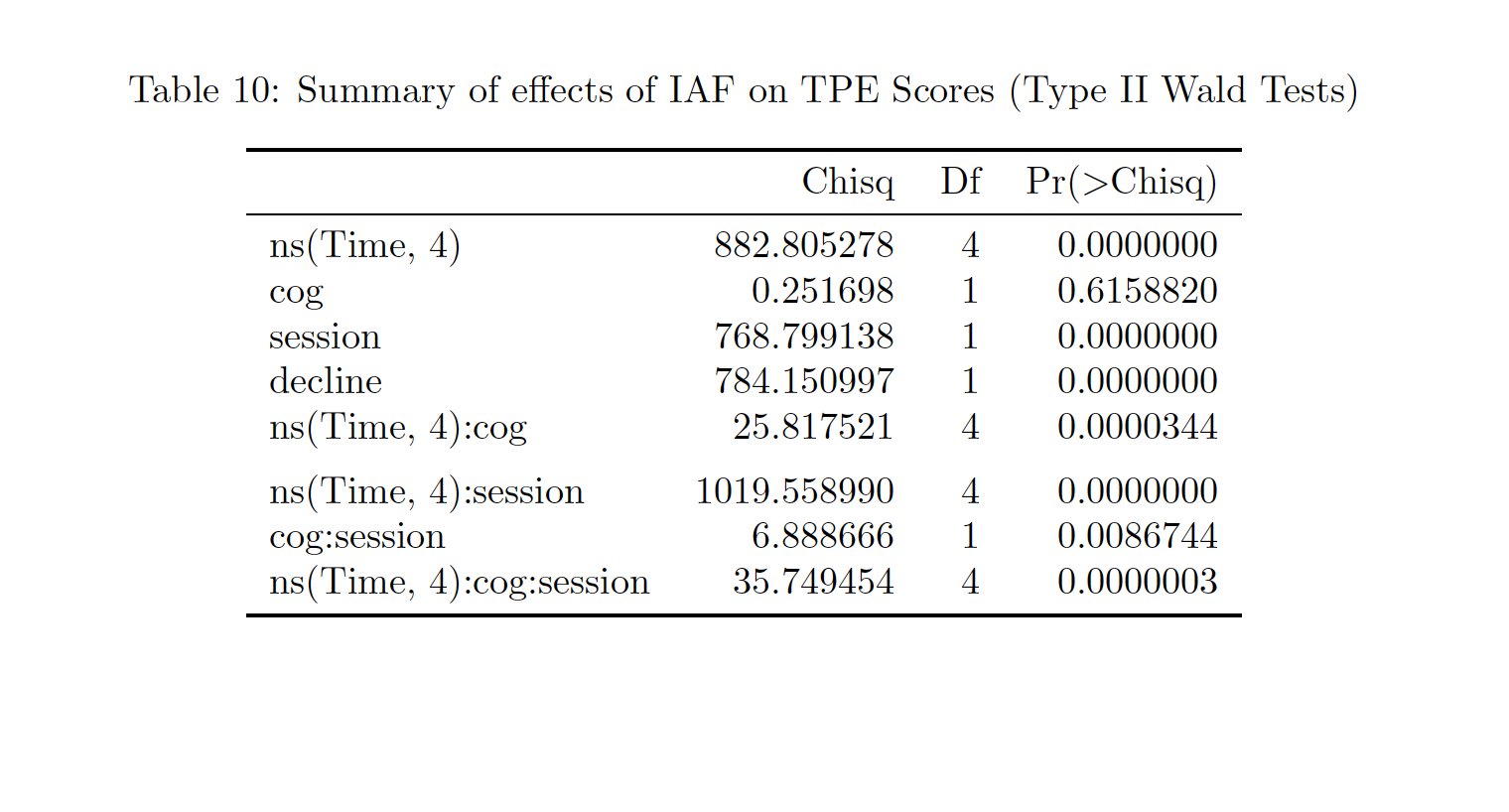

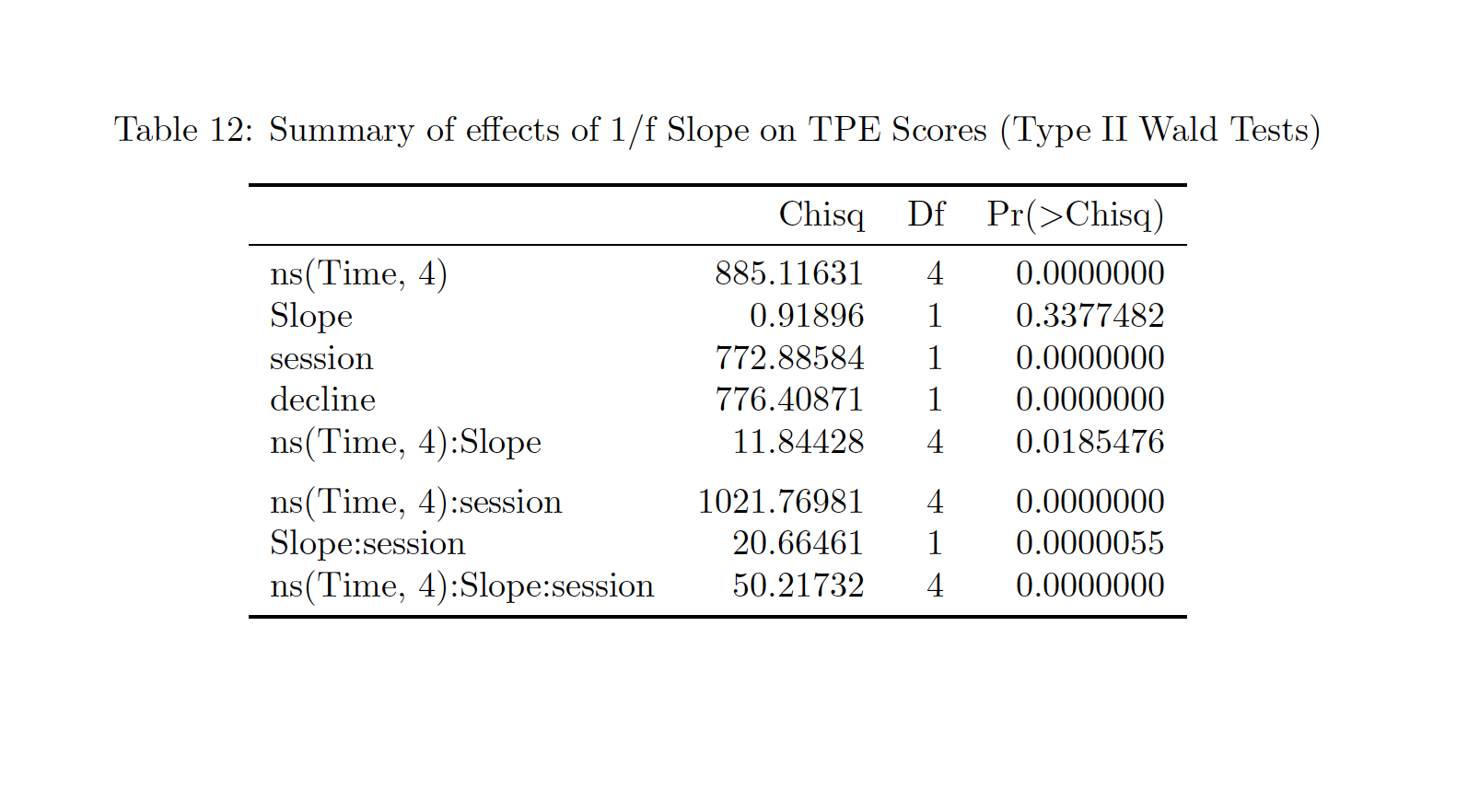

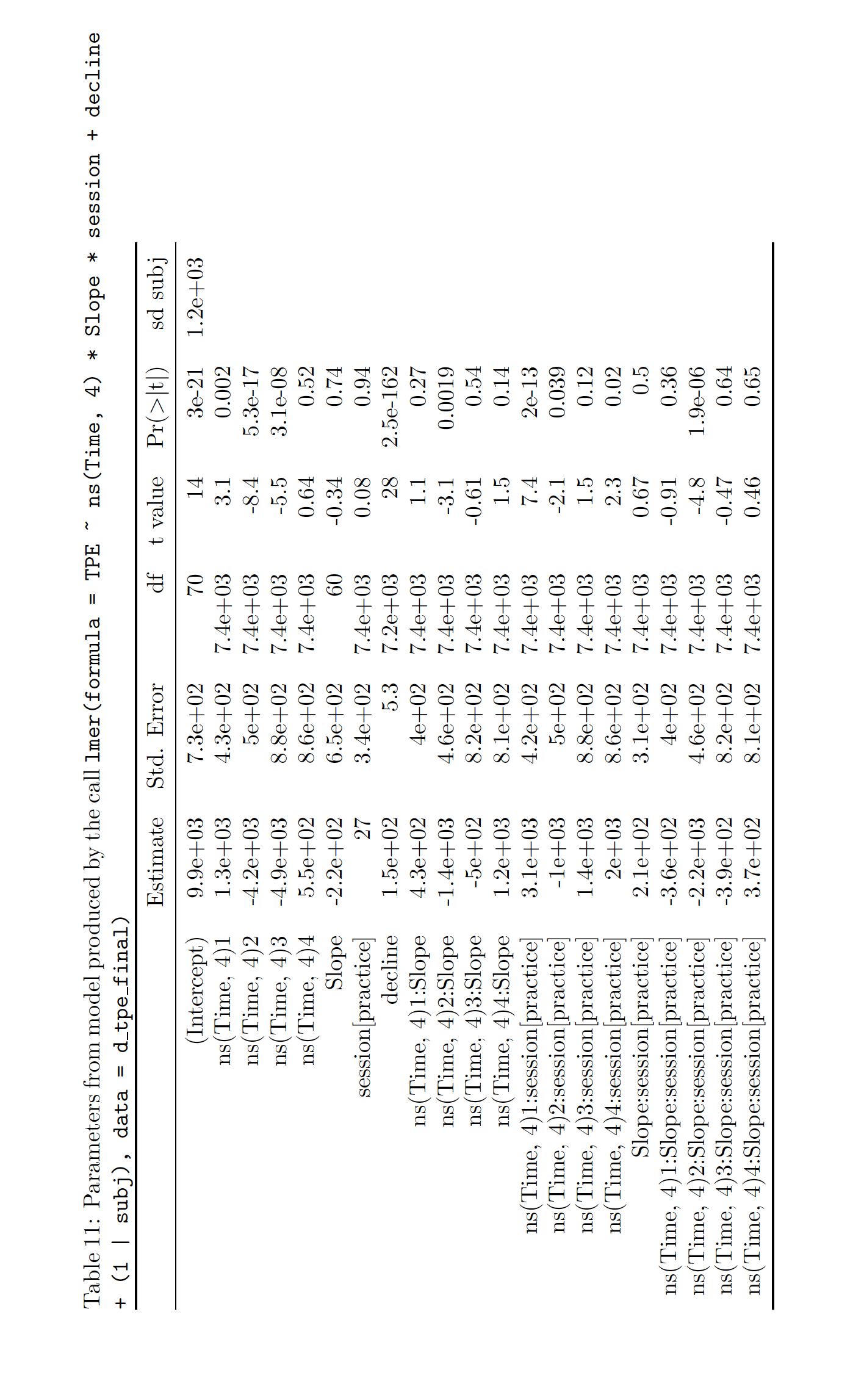

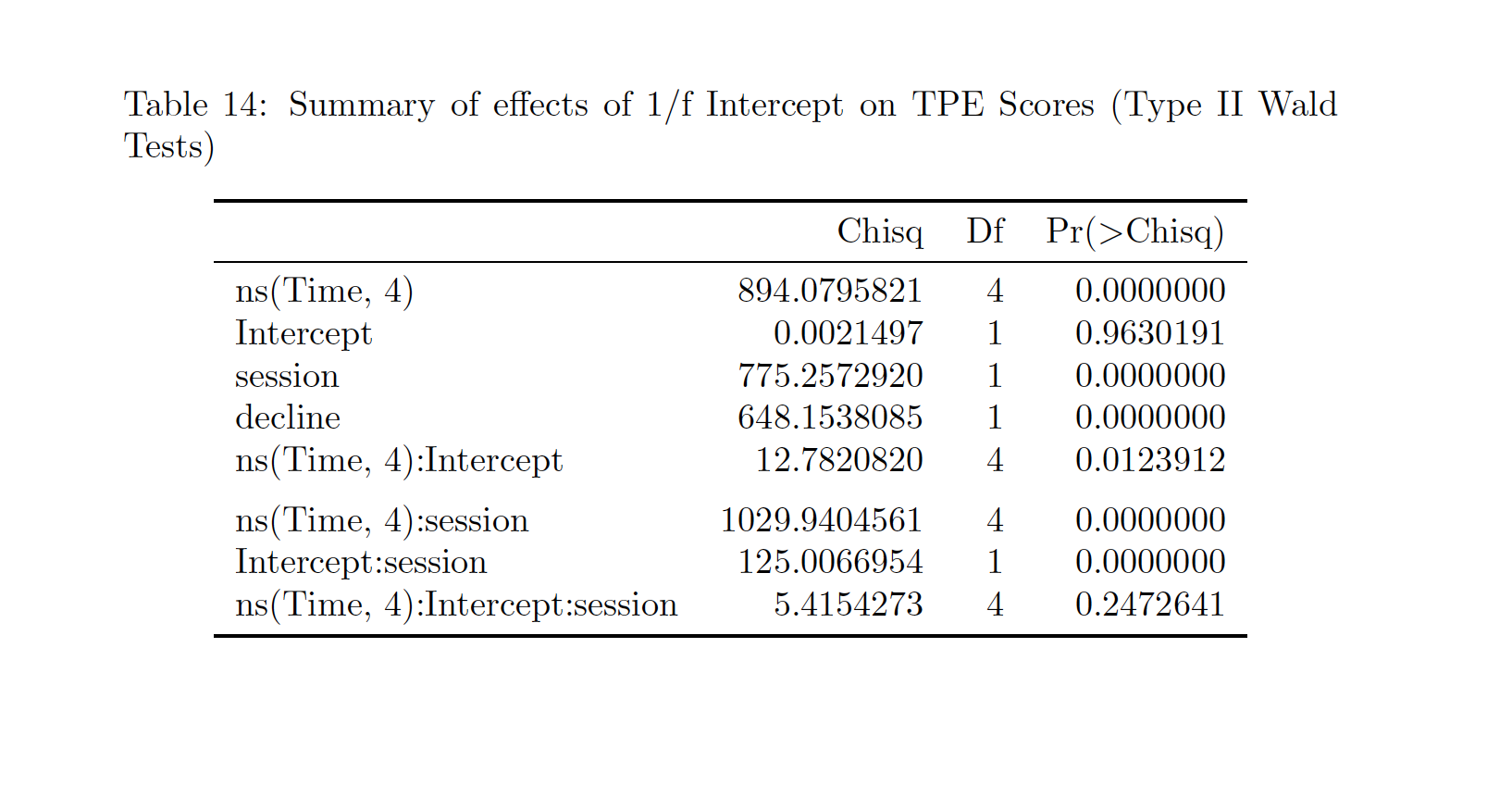

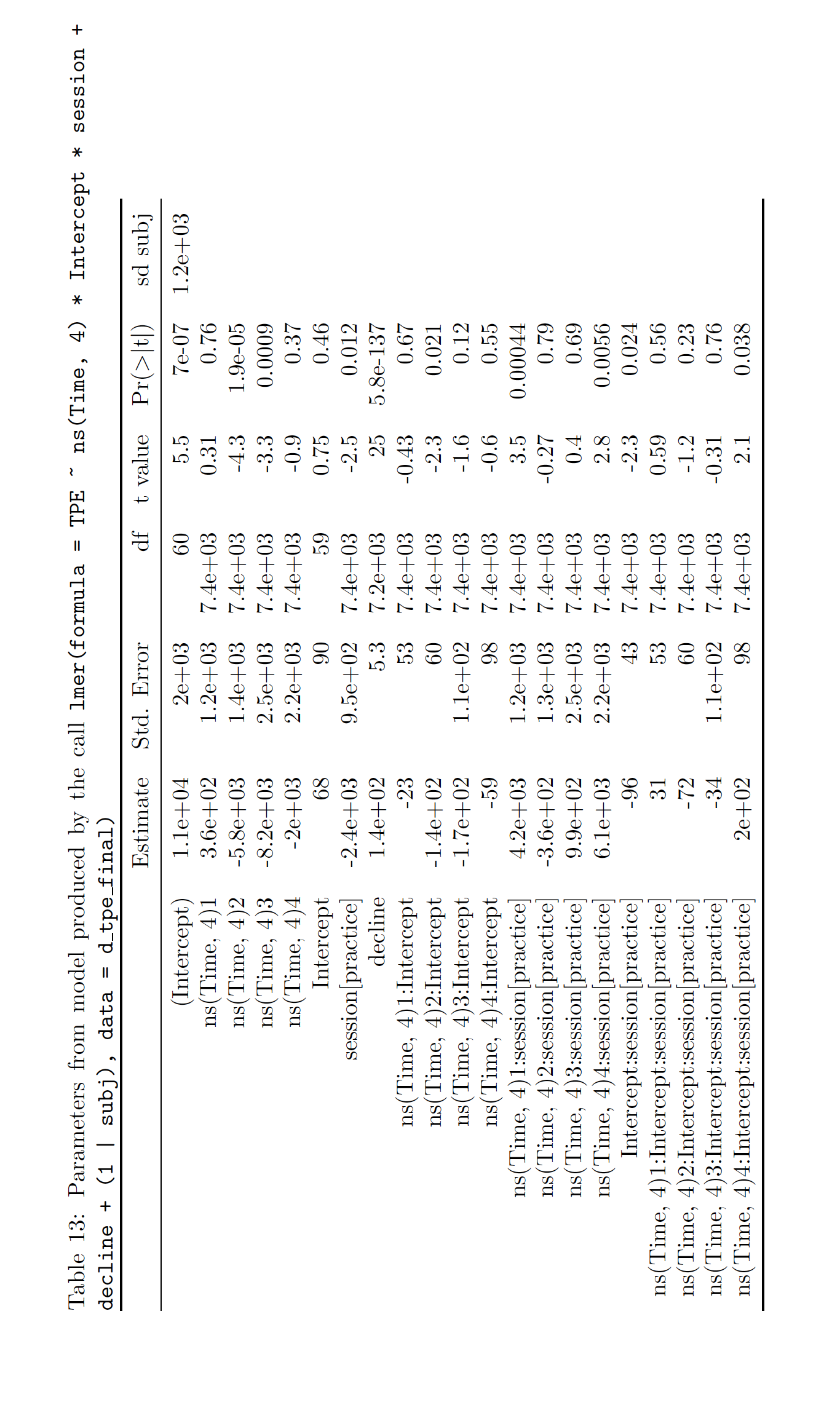
